# Resolving the oak tree of life: comparing RADseq and whole genome resequencing methods for oak phylogenetics

**DOI:** 10.64898/2026.05.14.725274

**Authors:** Andrew L. Hipp, Kieran N. Althaus, Elanor L. Fuller, Marlene Hahn, Drew A. Larson, Rebekah A. Mohn, Baosheng Wang, Paul S. Manos

**Author notes:** These authors contributed equally; all middle authors alphabetical by last name (Andrew L. Hipp, Kieran N. Althaus).

## Abstract

Forest trees pose numerous potential challenges to phylogenomic inference. Their large effective population sizes and relatively long generation times lead to deep allele coalescence and consequently incomplete lineage sorting (ILS), which biases inferences of divergence times toward older ages and introduces gene tree discordance. Deep phylogenetic divergences, reaching back into the Paleocene, introduce reference-mapping biases. Introgression—the movement of genes between lineages—may result in different phylogenies being inferred depending on which individuals are included in analysis, even if the plurality of the genome favors the divergence history unaffected by introgression. These factors influence phylogenetic inference across the Tree of Life but are particularly prevalent in forest trees. Oaks (*Quercus*) are notable for all three influences. In addition, our knowledge of the oak phylogeny is currently based strongly on restriction site associated DNA sequencing (RADseq) datasets published over the past decade, which may introduce additional sources of uncertainty. In this chapter, we analyze a 322-species RADseq dataset and genome resequencing data from across the genus to address sources of uncertainty in our understanding of the global oak phylogeny, which we hope will serve as a model for other research groups working on comparable woody plant groups.

## Introduction

Our ability to conserve and manage tree diversity rests on a clear understanding of what the species are, how they have evolved and may evolve in the future, and how species are adapted and interact. These three subjects—speciation, phylogeny, and adaptation—are particularly challenging to study in trees, whose large effective population sizes and high mean reproductive ages (Petit & Hampe, 2006; Daïnou *et al*., 2014) tend to result in higher rates of incomplete lineage sorting (ILS) than in shorter-lived perennial plants (Althaus & Hipp, 2026; Smith *et al*., 2026). High rates of interspecific hybridization and introgression also complicate tree phylogenetic history and may dramatically influence genetic variation within species (Suarez-Gonzalez *et al*., 2018). Thus, the life history of long-lived, outcrossing trees makes it particularly difficult to assess the boundaries of tree species and the relationships among and within forest tree populations.

Oaks (*Quercus* L., Fagaceae) are a model clade for understanding the evolution and ecology of forest tree diversity (Cavender-Bares, 2019; Kremer & Hipp, 2020). The first DNA-based studies of oak evolutionary history (Manos *et al*., 1999; Cavender-Bares *et al*., 2004; Oh & Manos, 2008; Denk & Grimm, 2010) analyzed relatively small numbers of genes or genome regions and converged on a set of broad relationships within *Quercus* (with formal names in parentheses following the phylogenomic classification of Denk *et al*., 2017):

- **Monophyly of the predominantly-American oak clade (*Quercus* subgenus *Quercus*)** comprising three major clades: the white oaks (sect. *Quercus*), the red oaks (sect. *Lobatae*), and the intermediate oaks (sect. *Protobalanus*); the southern live oaks (sect. *Virentes*) and deer oaks (sect. *Ponticae*) did not form clades separate from *Q.* sect. *Quercus* in these analyses.
- **Monophyly of the predominantly-Eurasian oak clade (*Quercus* subg. *Cerris*)** comprising two or three clades: the ring-cupped oaks (sect. *Cyclobalanopsis*) on one hand, the cork oaks (sect. *Cerris*) and the holly oaks (sect. *Ilex*) on the other (as either one or two clades, depending on dataset and analysis).

These early studies were unable to resolve relationships close to the tips of the phylogeny but raised hopes that there might be other, as-yet-undiscovered regions of the genome that might serve as a useful single marker for oak phylogenetics. A subsequent study by Hubert et al. (2014) used six genes selected from an estimated 621 candidate genes for potentially adaptive traits and 87 genes randomly selected from an oak EST (expressed sequence tag) database, an outstanding genomic resource at the time (Kremer *et al*., 2012; Plomion *et al*., 2016). Hubert’s work confirmed earlier results and provided additional support for separation of sections Ilex and Cerris, but relationships among close relatives remained elusive. Thus it seemed unlikely a single small set of genes would resolve phylogenetic relationships within subgenera.

A study published around the same time using amplified length polymorphisms (AFLPs; Pearse & Hipp, 2009) provided higher resolution near the tips of the tree, resolving relationships among closely related species and separating out previously elusive clades (e.g., distinguishing the Eurasian white oaks from the remaining oaks, and separating the Mexican red and white oaks as clades within their respective sections). AFLP bands sample for restriction site presence and absence across the genome, but they are anonymous without laborious cloning and sequencing. Moreover, bands of the same length for a given combination of restriction enzymes and selective nucleotides may come from entirely different regions of the genome, conflating homology and analogy and thus introducing noise into the data. The fact that these data yielded a hierarchical signal that accorded with expectations of the oak phylogeny suggested that the solution to resolving the oak phylogeny might lie not in a few loci each of high information content, but in many loci, whether their individual information content was high or, as in the AFLP dataset, low.

The advent of restriction-site associated DNA sequencing (RADseq; Baird *et al*., 2008) provided an economical balance between the analytical benefits of analyzing sequence data directly and the wide genome representation of AFLPs. There are many RADseq methodological variants, but all are reduced representation genome sequencing methods that sequence upstream and downstream from restriction sites. Fragments are then size-selected for sequencing, either after random shearing or a second digestion step (Andrews *et al*., 2016). The resulting datasets are large (often millions to tens of millions of basepairs) but generally sparse, with as much as 95% missing data (reviewed in Ree & Hipp, 2015).

RADseq has helped to resolve the evolutionary history of the oaks of North America and Mexico (Hipp *et al*., 2014, 2018, 2023; Cavender-Bares *et al*., 2015; Hauser *et al*., 2017; Fitz-Gibbon *et al*., 2017; Althaus *et al*., 2026), East Asia (Deng *et al*., 2018; Jiang *et al*., 2019), Europe (Denk *et al*., 2023), and the world (Denk *et al*., 2017; Hipp *et al*., 2020). RADseq has been useful for identifying the mosaic history of introgression within oaks (Eaton *et al*., 2015; Pham *et al*., 2017; McVay *et al*., 2017b,b; Ortego *et al*., 2018; Kim *et al*., 2018; Hipp *et al*., 2020) and identifying minimum numbers of SNPs that suffice to distinguish species (Fitzek *et al*., 2018; Hipp *et al*., 2019). RADseq data (previous citations) have provided a robust framework phylogeny for the oaks (Fig. 1) that is also supported by subsequent genome resequencing and sequence-capture datasets (Zhou *et al*., 2022; Yang *et al*., 2023; Liu *et al*., 2025; Shen *et al*., 2025).

**Figure 1.**
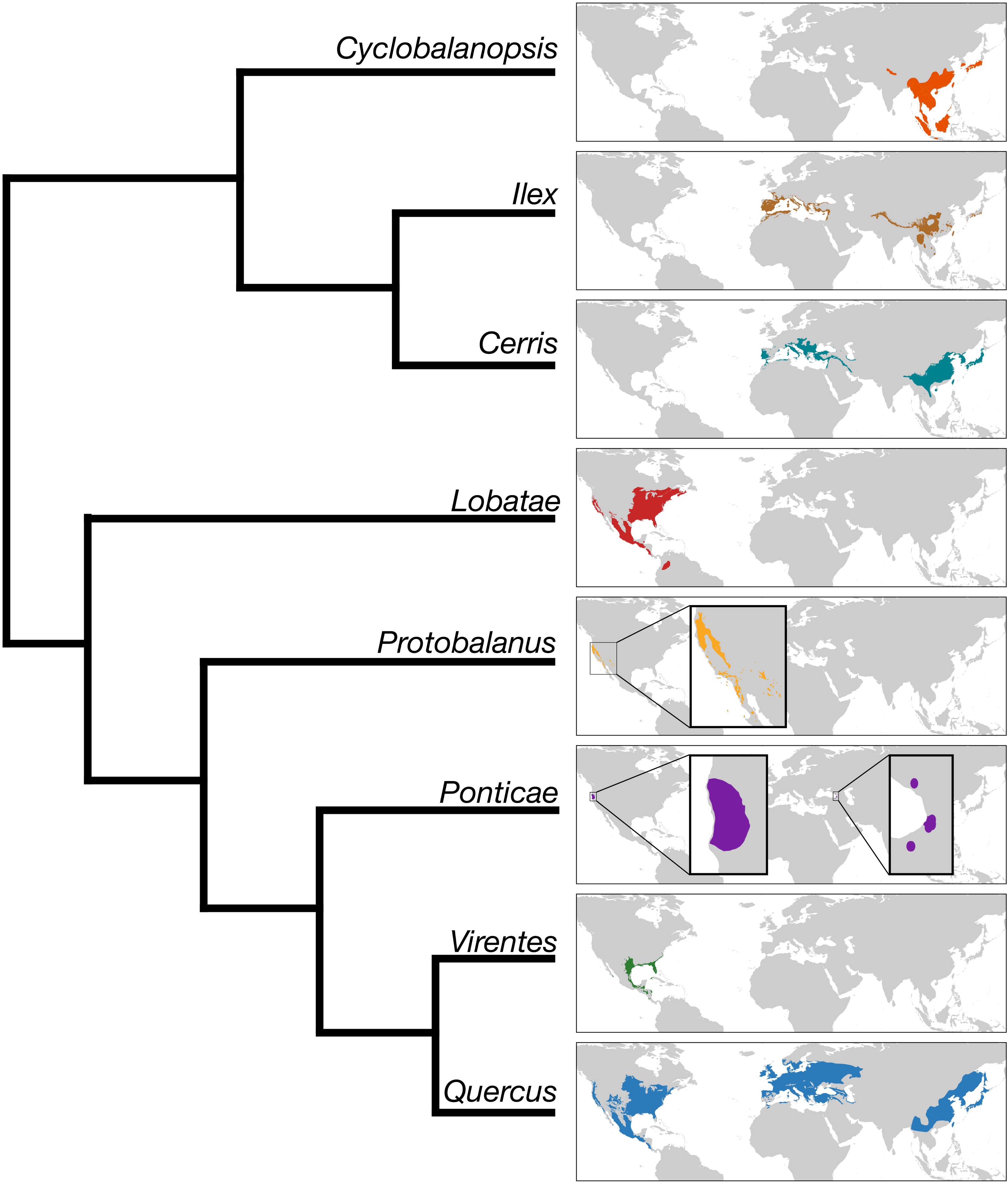
Framework oak phylogeny with section maps. Framework oak phylogeny follows Hipp et al. 2018, 2020. Range maps of the 8 named clades (sections) are from Althaus and Martín Sánchez (in Hipp, 2024, pp. 126–7).

While RADseq data have contributed a great deal to our understanding of oak evolutionary history, they have a few inherent limitations. First, because RADseq data are based on short-read sequencing, RADseq locus homology assessment can be problematic. This is probably not a great problem in contemporary oaks, where polyploidy is rare (though see Hu *et al*., 2026) and genome structure is relatively conserved (Kapoor *et al*., 2023; Larson *et al*., 2025; Aközbek *et al*., 2026). The use of short read data not pinned to particular target loci introduces a second problem: each locus provides little information about relationships, making it difficult to tease apart ILS and hybridization histories. Third, RADseq data are generated, for the most part, one short read upstream and downstream from restriction sites. This makes them genomically sparse. As a consequence, RADseq data are less useful for genomic window analysis and connecting SNP variation to function (Lowry *et al*., 2017). The advent of sequence capture probesets for *Quercus* (Crowl *et al*., 2020; Morales-Saldaña *et al*., 2024) and comparative resequencing data (Larson *et al*., 2025; Mohn *et al*., 2026) offer increasingly affordable and scalable options for oak phylogenetics.

In this chapter, we ask three questions about the reliability of RADseq phylogenies: (1) Do RADseq data return phylogenies that differ systematically from SNP datasets derived directly from genomic data? (2) Does the choice of a particular reference genome for calling genomic variants (SNPs) influence phylogenetic inferences in genomic and RADseq data? and (3) Is variation among and within phylogenomic datasets at least partially attributable to gene flow? To address these questions, we analyze a large (322-species) RADseq dataset for *Quercus* using seven reference genomes plus *de novo* clustering; a SNP set generated from a collection of 76 individuals with whole genome resequencing data mapped to reference genomes; and a simulated RADseq dataset generated from the 76-individual genomic data. We assess phylogenetic variation among and within datasets and investigate correlations between sources of variation and histories of introgression. In the end, our exploration of these data sources aims to expose the strengths and limitations of the RADseq approach in phylogenetics when evaluated against whole genome resequencing, as a guide to future data use and reuse.

## Methods

### RADseq data generation and clustering

RADseq data for 322 samples were previously sequenced on either the Illumina Genome Analyzer IIx, Illumina HiSeq 2000, HiSeq 2500, HiSeq4000 or NovaSeq 6000 (Cavender-Bares *et al*., 2015; Fitz-Gibbon *et al*., 2017; McVay *et al*., 2017b,a; Deng *et al*., 2018; Hipp *et al*., 2018, 2020; Jiang *et al*., 2019; Althaus *et al*., 2026) (Table S1). Seven reference genomes from different sections of the genus *Quercus* were downloaded to use as reference genomes: *Quercus glauca* from section *Cyclobalanopis* (Luo *et al*., 2024), *Q. variablis* from sect. *Cerris* (Wang *et al*., 2023), *Q. longispica* from sect. *Ilex* (Ren *et al*., 2025), *Q. rubra* from sect. *Lobatae* (Kapoor *et al*., 2023), *Q. tomentella* from sect. *Protobalanus* (Mead *et al*., 2024), *Q. virginiana* from sect. *Virentes* (Aközbek *et al*., 2026), and *Q. alba* from sect. *Quercus* (Larson *et al*., 2025).

ipyrad v.0.9.81 (Eaton & Overcast, 2020) was used to filter out all sequences that had more than five bases of quality score <20, and to remove adapter/primers. All sequences were trimmed to 90 bps. For the *de novo* clustered dataset, sequences were clustered at 85% sequence similarity; for datasets mapped to separate reference genomes (rather than clustered *de novo*), ipyrad was branched off at step three to map reads to each of the respective genomes instead of clustering. Parameters for all runs were set so that there was a minimum of six sequence reads per individual per locus. Loci were excluded if they had a cluster depth of greater than 10,000 sequences. Loci were excluded from the dataset if they had more than two alleles, 5% Ns, or 5% heterozygotic positions in the consensus sequence. All final alignment datasets were filtered to contain loci that were present in at least four individuals.

Throughout the chapter, these datasets and the trees inferred from them are referred to using the term “empirical RADseq”.

### Whole genome resequencing data and pseudo-reference genomes

#### Sampling resequencing data and generating pseudo-references

Genome-wide SNPs were called for 76 species, covering the taxonomic breadth of Quercus and four *Lithocarpus* species as an outgroup (Table S2). Reads included novel sequence data and reads from the Sequence Read Archive (SRA). Paired end sequences were downloaded from the SRA with fasterq-dump v3.0.5 (SRA Toolkit Development Team, 2026). Reads of one individual were trimmed to remove over-represented sequences with Skewer v0.2.2 (Jiang *et al*., 2014); parameters for this step are available in the accompanying repository submission. Then, reads for all samples were trimmed using fastp v0.23.4 (Chen *et al*., 2018) and the options "--low_complexity_filter --complexity_threshold 25” --trim_poly_g". This and most other steps were completed using GNU-parallel (Tange, 2023). Prior to trimming with fastp, All reads were aligned with bwa mem (Li & Durbin, 2009) to two different reference genomes, that of *Quercus alba* (Larson *et al*., 2025) and that of *Quercus variabilis* (Wang *et al*., 2023) and all downstream steps were conducted for each sample/reference combination. The resulting sam files were sorted using the sort command from SAMtools v1.17 and the “--write-index” option (Li *et al*., 2009). Duplicate reads were marked using the MarkDuplicates command in picard v2.27.3 (‘Picard Tools - By Broad Institute’). Coverage was calculated for each base using the Depth command in SAMtools and the option “-aa”. A pileup was generated for each set of reads using the mpileup command from BCFtools v1.17 (Li, 2011) and the options “-Ou -a FORMAT/AD,FORMAT/DP --skip-indels --min-BQ 15 --min-MQ 15” specified followed by SNP calling with the call command and options “-m --gvcf 10 -Oz”. The BCFtools command filter, with options "--set-GTs. -e ’FORMAT/DP < 8’ -Oz", was used to code all genotypes with less than 8× coverage as missing data and the command index was used to generate an index for each VCF. A pseudo-reference sequence was generated for each sample and reference combination using the consensus command in BCFtools with the options "--haplotype I --absent ’?’ --missing ’n’ --mark-del ’-’". VCFs for all samples were then merged into a single VCF per reference genome using the merge command in BCFtools.

Throughout the chapter, these datasets and trees inferred from them are referred to using the terms “whole genome resequencing” or “resequencing”.

#### Simulating RADseq data from pseudo-references

For each of the 76 samples for which we generated a pseudo-reference sequence, we extracted restriction site loci using RADinitio (Rivera-Colón *et al*., 2021). Using a whole genome reference sequence, RADinitio simulates genomic data and different stages of the RADseq library preparation, from population sampling to library enrichment. RADinitio simulates the various sources of error associated with RAD sequencing, including as locus-sampling and PCR errors. Because RADinitio requires haploid genomes without gaps to conduct its simulations, we randomly resolved heterozygous sites with the program seqtk v 1.5-r133 (Li, 2025) and removed all gapped sites with a custom python script before simulation. Following the library preparation parameters outlined in Hipp et al. (2014), we simulated a standard RADseq library with a mean insert size of 450 bp and a standard deviation of 50 bp. We simulated 12 PCR cycles, a coverage of 30× and a read length of 100 bp. All other library preparation parameters were set to the RADinitio default values, including (among other defaults): PCR Depth/Complexity ratio = 0.1; Probability of indels = 0.01, with insertions and deletions equal rates; and probability of PCR errors = 4.4e-7. The result of our simulation runs are RADloci from each of the 76 resequencing samples, with simulated error introduced by RAD sequencing. Reads for RADloci were output as FASTQ files. Code for simulations is cached at https://doi.org/10.5281/zenodo.20084868 (v0.9-7, 2026-05-14).

FASTQ files were assembled into loci for phylogenetics using ipyrad v.0.9.81 (Eaton & Overcast, 2020), following the same clustering and assembly approaches used with the empirical RADseq data (see methods above). We ran assembly steps mapping to the *Quercus alba* reference genome for the pseudo-references that were generated by mapping to *Q. alba* (Larson et al. 2025); a second mapping to the *Quercus variabilis* genome (Wang *et al*., 2023) for the pseudo-references that were generated by mapping to *Q. variabilis*; and using de novo clustering for both sets of simulated RADseq datasets. This yielded a total of four simulated RADseq datasets. We filtered loci to only those present in at least four individuals.

Throughout the chapter, these datasets and trees inferred from them are referred to using the terms “simRAD” or “simulated RADseq”.

### Phylogeny estimation

#### RADseq datasets (empirical and simulated)

The most comprehensive oak phylogenies to date (Hipp *et al*., 2020; Althaus *et al*., 2026) were generated using maximum likelihood (ML) conducted with RAxML v8.2.4 (Stamatakis, 2014). For consistency, we likewise used RAxML in our analysis of our data, running ML analyses of concatenated data for all phylogenetic estimates. We estimated one phylogeny for each reference genome dataset, generating seven maximum likelihood estimates. Each genome alignment was subset to keep only sites with data in at least 60% of individuals with the *pxclsq* command in phyx v1.1 (Brown *et al*., 2017). For the simulated RADseq data, we estimated a phylogeny for data mapped to both the *Q. alba* and *Q. variabilis* genomes. Both sets of analyses were run with a GTR+G model of nucleotide evolution and 200 bootstraps to evaluate support for the maximum likelihood topology.

#### Whole genome resequencing datasets

Given the size of the whole genome resequencing datasets, we opted to subset the VCF for each alignment and analyze SNPs only. We filtered our VCF to include SNPs only with BCFtools (Danecek *et al*., 2021) and used vcf2phylip v2.0 (Ortiz, 2019) to convert the VCF into PHYLIP format. We refer to these as the “reseq datasets”. We estimated two phylogenies—one for the dataset generated using the *Q. alba* reference, one for the dataset generated using the *Q. variabilis* reference—using the GTR model with ascertainment bias and 200 bootstrap replicates in RAxML v8.2.4. We refer to these as the “reseq trees”.

To estimate introgression in the whole genome resequencing datasets, we calculated f-branch statistics (Malinsky *et al*., 2018) with Dsuite v0.5 r58 (Malinsky *et al*., 2021), the full combined VCF for each reference, and the inferred species tree with *Lithocarpus* as the outgroup. The f-branch results were visualized using utilities included with Dsuite. To assess introgression signals among oak sections, a second f-branch analysis was conducted in which all species of each section were treated as the unit of analysis (*i.e.*, each section was treated as a population). Recent work evaluating the power of *D*- and *f-* statistics to detect hybridization and introgression events has highlighted the impact of even slight variation on the rate of molecular evolution (Frankel & Ané, 2023; Pang *et al*., 2025). To evaluate molecular rate variation across our genomic dataset, we followed Pang *et al*. (2025) and conducted the relative-rate test (Graur & Li, 2000) in non-overlapping sliding windows across our *Q. alba*-mapped VCF file. We measured the genetic distance between each pair of species in our dataset using the JC69 model of molecular evolution on each 50-kb genomic window in our alignment. Relative rates were calculated by comparison with *Lithocarpus*, our outgroup. See Supplementary File 1 of Pang *et al*. (2025) for a more detailed description of the method, and the repository accompanying this chapter for analysis scripts. The results are summarized at the section-level, but the full, species-level, output is available in the supplement.

#### Topology comparisons

To visualize variation among datasets and analyses, all topologies were pruned down to 59 tips representing species sampled both in the empirical RADseq and the reference genomes dataset (and thus present also in the simulated RADseq dataset), as well as the empirical RADseq tree published in Hipp et al. (2020). This is somewhat unfortunate, as our empirical RADseq data includes more than 300 species, but it focuses analysis on some of the key relationships among and within larger clades. Pairwise entropy distances between all trees were calculated using the clustering information distance in TreeDist v. 2.11.1 (Smith, 2020a,b, 2022) in R v. 4.5.2 (R Core Team, 2025). Results were visualized using classic (metric) multidimensional scaling (cmdscale) in core R, with *K* = 2 dimensions for ease of visualization, and plotted in ggplot2 v. 4.0.1 (Wickham, 2016). Tree ordination groups were identified visually. Pairwise comparisons between selected trees were visualized using *cophylo* in phytools v. 2.5.2 (Revell, 2024).

To identify clades most strongly influencing distribution of trees in treespace, each tree (including ML and bootstrap trees) was coded for monophyly of six groups observed in our analysis and known from previous work to resolve variously in different phylogenomic analyses: the California white oaks; the Mexican white oaks; the Mexican white oaks + the eastern North American subsection *Stellatae*; the Eurasian Roburoid white oaks + the eastern North American *Albae*; *Quercus macrocarpa* + *Q. bicolor*; and *Q. macrocarpa* + *Q. bicolor* + *Q. lobata* (McVay *et al*., 2017b,a; Crowl *et al*., 2020; Hipp *et al*., 2020; Zhou *et al*., 2022; Larson *et al*., 2025); subsections here and throughout the chapter follow Manos and Hipp (2021). Trees possessing each clade were assigned a value of 1, those which lacked it were assigned a value of 0. These values were then projected onto ordinations using the envfit function of vegan v 2.7-2 (Oksanen *et al*., 2025).

#### Code availability

Code to execute analyses in this chapter is archived at https://doi.org/10.5281/zenodo.20084868 (v0.9-7, 2026-05-14).

## Results

### Datasets

Empirical RADseq statistics are reported in previous papers (Hipp *et al*., 2020; Althaus *et al*., 2026), and clustering statistics for the current study are available in the online supplement (https://doi.org/10.5281/zenodo.20084868, subfolder /topologies/data/ipyrad-stats/empirical). Total dataset filtered loci using reference-guided clustering range from 115,998 with the *Q. rubra* reference to 123,458 with the *Q. longispica* reference; *de novo* clustering yielded 153,489 loci, 24% more than the highest number recovered using reference-guided clustering. Average number of filtered loci per individual follow the same rank order: 7,903 +/- 6,397 (sd) for the *Q. rubra* reference-mapping dataset, 7,924 +/- 6,790 for the *Q. longispica* reference-mapping dataset, 8,128 +/- 2,404 for the *de novo* clustering dataset, an increase of only 2.57% over the *Q. longispica* reference dataset. This is a wide range of variation within each dataset; excluding one poorly sequenced individual with less than 1000 loci (*Quercus phanera*), the *Q. rubra*-mapped empirical RADseq dataset exhibits a 7.0-fold difference between the individual with fewest sampled loci (1,864 in *Quercus cedrosensis*) and the largest (13,107 in *Quercus microphylla*, which included results of two sequencing runs as one was of very low quality).

Pseudo-references for 76 samples were generated by mapping 65 published and 11 unpublished resequencing datasets (Table S2) to both the *Q. alba* and *Q. variabilis* reference genomes. Missing data per genome under this method ranged from 18.2% to 72.6% (47.0 +/- 11.8 sd) for the *Q. alba* reference and 21.3% to 68.6% (54.0 +/- 10.2 sd) for the *Q. variabilis* dataset. The VCF files, filtered to include only variable sites for each reference genome, included 313.97 million and 303.12 million SNPs for the *Q. alba* and *Q. variabilis* aligned data, respectively (summary files in online supplement: https://doi.org/10.5281/zenodo.20084868, subfolder /topologies/data/pseudoref-stats). After filtering to sites with allele count (AC) > 1 in called genotypes, we retained 92 million sites in the *Q. alba* reference and 85 million in the *Q. variabilis* reference. Both sets of VCFs are highly concordant in other genotype properties, such as ts/tv ratio, fraction of multiallelic sites, singletons fractions and substitution-type rank order.

Simulated RADseq reads from the resulting pseudo-reference genome sequences were assembled in ipyrad under four conditions: *de novo* and reference-mapped for both the *Q. alba* and *Q. variabilis-*mapped pseudo-reference genome sets, using the corresponding reference for ipyrad assembly (ipyrad assembly and clustering statistics: https://doi.org/10.5281/zenodo.20084868, subfolder /topologies/data/ipyrad-stats/simulated). The choice of reference genome had much larger effect on locus yield than the choice of assembly method: *Q. alba-*referenced datasets retained 11,432 loci *de novo* and 11,210 loci with reference mapping, whereas *Q. variabilis*-referenced datasets retained 6,395 and 6,278, respectively (Table S3). Within each reference, *de novo* and reference-mapped assemblies produced nearly identical locus counts (within 2%), while locus retention was overwhelmingly limited by the minimum-sample filter among all assembly methods (> 97% of all filtered loci in every assembly). Reference mapping recovered longer loci and more variation than *de novo* clustering: concatenated sequence matrices were 60% larger, and SNP matrices contained 30% more variable and parsimony-informative sites. Missing data were higher in reference-mapped matrices than in the *de novo* matrices.

The simRAD data matrices recovered many more loci for the *Quercus alba* - mapped pseudo-reference genome set (11,432 and 11,210 for *de novo* and reference-guided assembly respectively) than for the *Q. variabilis* - mapped reference-genome set (6,395 and 6,278 respectively; Table S3). However, the difference between *de novo* and reference-guided assemblies was negligible (1.9%; Table S4a). By contrast, the *de novo* empirical RADseq data matrix recovered 28.4% more loci than the reference-guided empirical RADseq data matrices (Table S4a). This is likely due in part to the fact that the reference-guided datasets concatenate loci that flank the same restriction site; thus, for example, while the average assembled sequence (‘locus’) length for the *de novo* empirical clustering dataset is 86.9 bp, the average assembled sequence (‘locus’) length for the *Q. alba* reference-guided dataset is 164.8 bp.

Datasets exhibited large differences in the number of unique phylogenetic site patterns. On average, the genome resequencing and simRAD data matrices respectively had 62× and 2.9× as many phylogenetic site patterns as the reference-guided empirical RADseq data matrices (Table S4b). The *de novo* clustered empirical RADseq dataset had 24× as many site patterns as the average of reference-guided empirical RADseq data matrices (Table S4b), which may reflect the noise of non-homologous loci clustering in the absence of a reference.

### Phylogenetic inferences from individual datasets

#### Maximum likelihood trees and bootstrap sets

The empirical RADseq topologies of 322 taxa (Fig. S1a–S1h) support all sections at 100% bootstrap support, with three notable exceptions. (1) The intermediate oaks, section *Protobalanus*, are supported at 47% in the RAD_variabilis_ tree (Fig. 2g, S1g) due to instability in the placement of *Q. cedrosensis* (which has the third-lowest locus coverage of all samples in our study, at an average of 1,820 loci across analyses, compared to the mean of 7,809 +/- 6,207 [sd]) for all samples). (2) The white oaks, section *Quercus*, are paraphyletic in the *de novo* clustering phylogeny, which has sect. *Virentes* as a subclade of sect. *Quercus* with 100% bootstrap support (Fig. 2a, S1a). (3) Three of the empirical RADseq datasets place *Notholithocarpus* within *Quercus* with high support (97–100%), rendering *Quercus* paraphyletic: RAD_glauca_ (Fig. S1c), RAD_rubra_ (Fig. S1e), and RAD_tomentella_ (Fig. S1f). Aside from these discrepancies, the relationships among sections are the same for all the empirical RADseq phylogenies (Fig. 2, S1).

**Figure 2.**
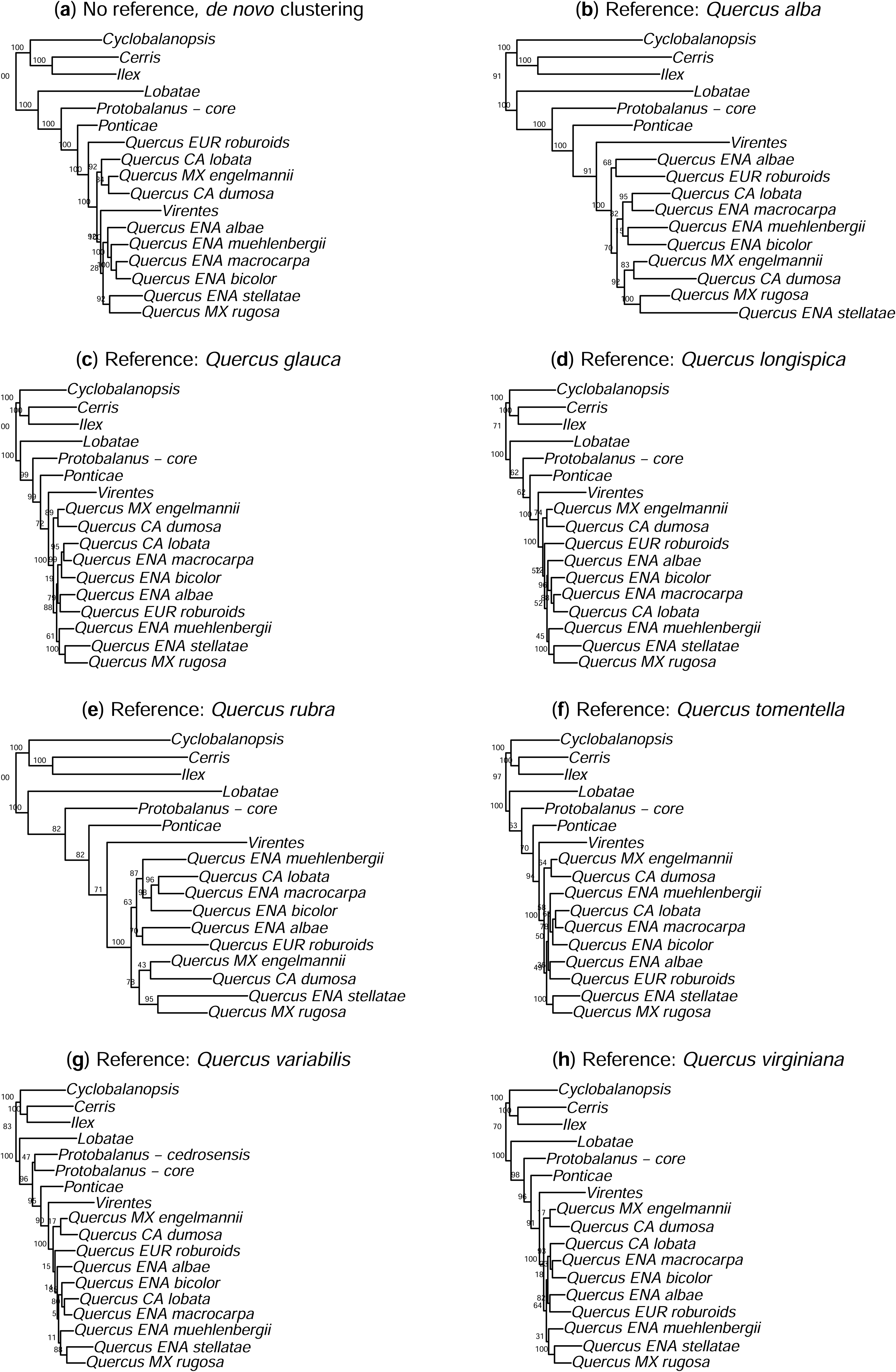
Empirical RADseq phylogenies, section summaries. Tree name subscripts indicate which of 7 reference genomes was used in ipyrad locus-calling, except for (a) *de novo* clustering (no reference). References used: (b) *Q. alba*, section *Quercus*; (c) *Q. glauca*, sect. *Cyclobalanopsis*; (3) *Q. longispica*, sect. *Ilex*; (e) *Q. rubra*, sect. *Lobatae*; (f) *Q. tomentella*, sect. *Protobalanus*; (g) *Q. variabilis*, sect. *Cerris*; (h) *Q. virginiana*, sect. *Virentes*. See text for citations.

The two whole genome resequencing datasets (Fig. 3, S2) recover virtually identical topologies, with bootstrap support at 100% for all nodes except for some variation within the Eurasian white oaks (bootstraps < 100% indicated on Fig. 3). There are two very minor topological differences between the trees based on mapping to the *Q. alba* reference (Fig. 3a, S2a) versus the *Q. variabilis* reference (Fig. 3b, S2b): the position of *Q. griffithii* (in the east Asian white oaks, sect. *Quercus*) and the resolution of *Q. fleuryi*, *Q. hui*, and *Q. edithiae* (sect. *Cyclobalanopsis*) (Fig. 3; conflicting nodes shown in red). Within sect. *Quercus* (the white oaks), both trees are in agreement with previous RADseq analyses (Hipp *et al*., 2018) in resolving the Mexican oaks (including *Q. engelmannii*) as monophyletic and sister to subsection *Stellatae*, and the Eurasian white oaks (the “Roburoids”) as sister to the three species of subsection *Albae*. In contrast with previously-published RADseq trees, they resolve the Californian white oak *Q. lobata* as sister to the trio of (*Q. macrocarpa*, (*Q. lyrata*, *Q. bicolor*)), embedded within subsection *Prinoideae* of section *Quercus*, rather than as sister to *Q. dumosa*, the other California white oak sampled in this study (excluding *Q. engelmannii*, which is a Mexican white oak by origin; see discussion below).

**Figure 3.**
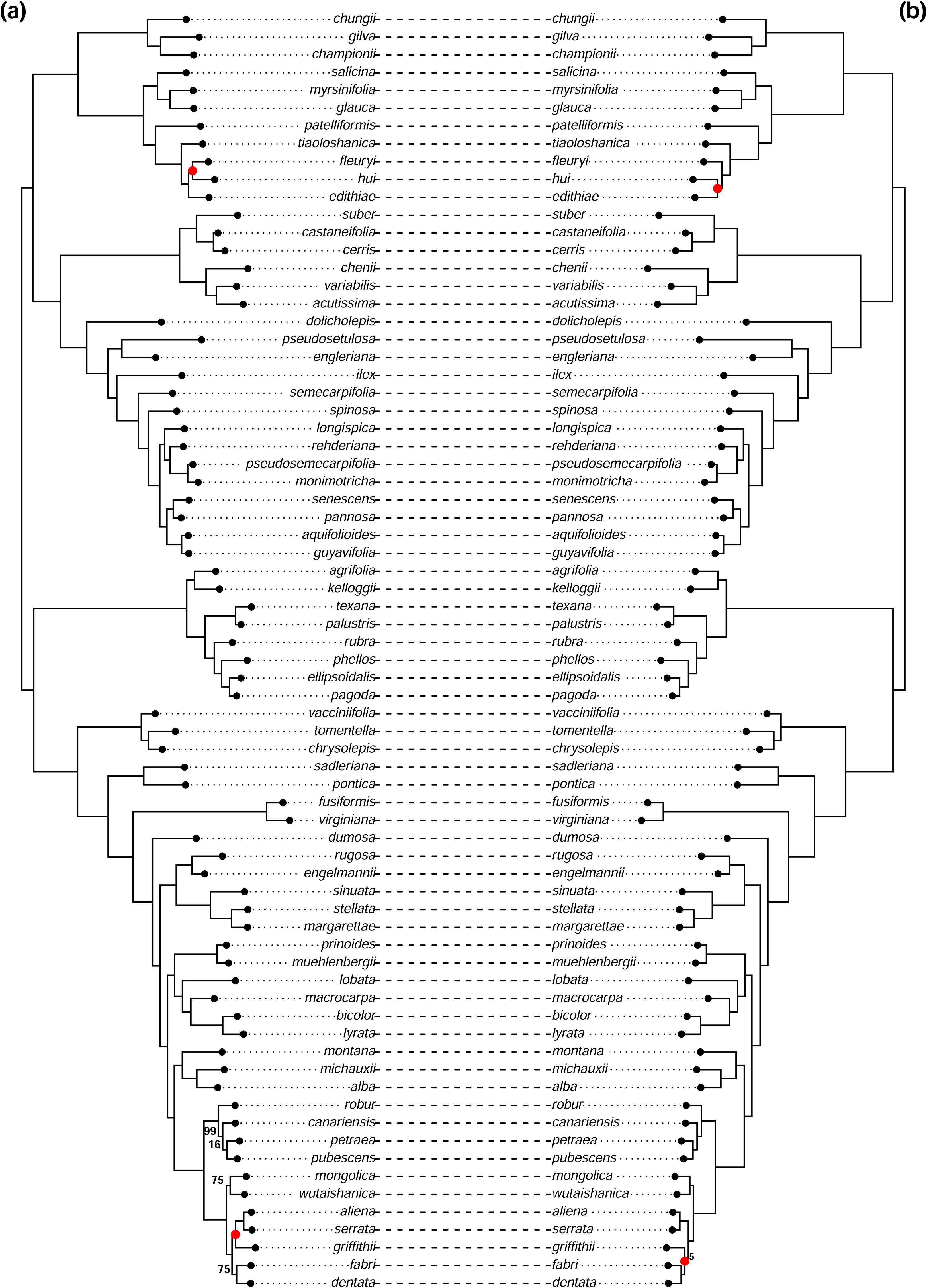
Whole genome resequencing (SNP) phylogenies, all ingroup taxa. Trees were estimated using RAxML on a SNP matrix from reference genomes and resequencing data for 72 ingroup taxa (shown) plus 4 *Lithocarpus* outgroups (not shown, but present in tree files in the online supplement). SNPs were called based on mapping of all genomes to either (a) the *Quercus alba* reference genome or (b) the *Q. variabilis* reference genome (citations in text). Red dots at two nodes on each panel highlight clades present in one tree but not the other.

Simulated RADseq (simRAD) trees have much lower bootstrap support within major clades than the reseq trees do, indicating a high degree of phylogenetic variance (Fig. S3a–d). While the whole genome resequencing data trees have mean bootstrap support of 98.1 (*Q. alba* reference) and 98.4 (*Q. variabilis* reference), the simulated RADseq datasets have a mean bootstrap support of 86.3 +/- 1.0 [sd], comparable to the empirical RADseq datasets (mean bootstrap = 84.1 +/- 3.2 sd).

Excluding the *de novo* clustering of RADseq data, trees pruned to the 59 taxa shared between all analyses—a total of 13 trees: 7 reference-guided empirical RADseq trees, 4 simulated RADseq trees, and 2 resequencing trees—unanimously support (1) monophyly of the eight oak sections recognized in Denk et al. (2017); (2) relationships among all sections; and (3) monophyly of the Eurasian or Roburoid white oaks, as well as reciprocal monophyly of the European and east Asian Roburoids (Fig. 4). Additionally (but not labelled in Fig. 4), the strict consensus of these 13 trees supports (4) reciprocal monophyly of the East Asian and Mediterranean clades of sect. *Cerris*; (5) reciprocal monophyly of two clades identified but not named in previous analyses of sect. *Cyclobalanopsis*; and (6) reciprocal monophyly of the East Asian and European Roburoid clades.

**Figure 4.**
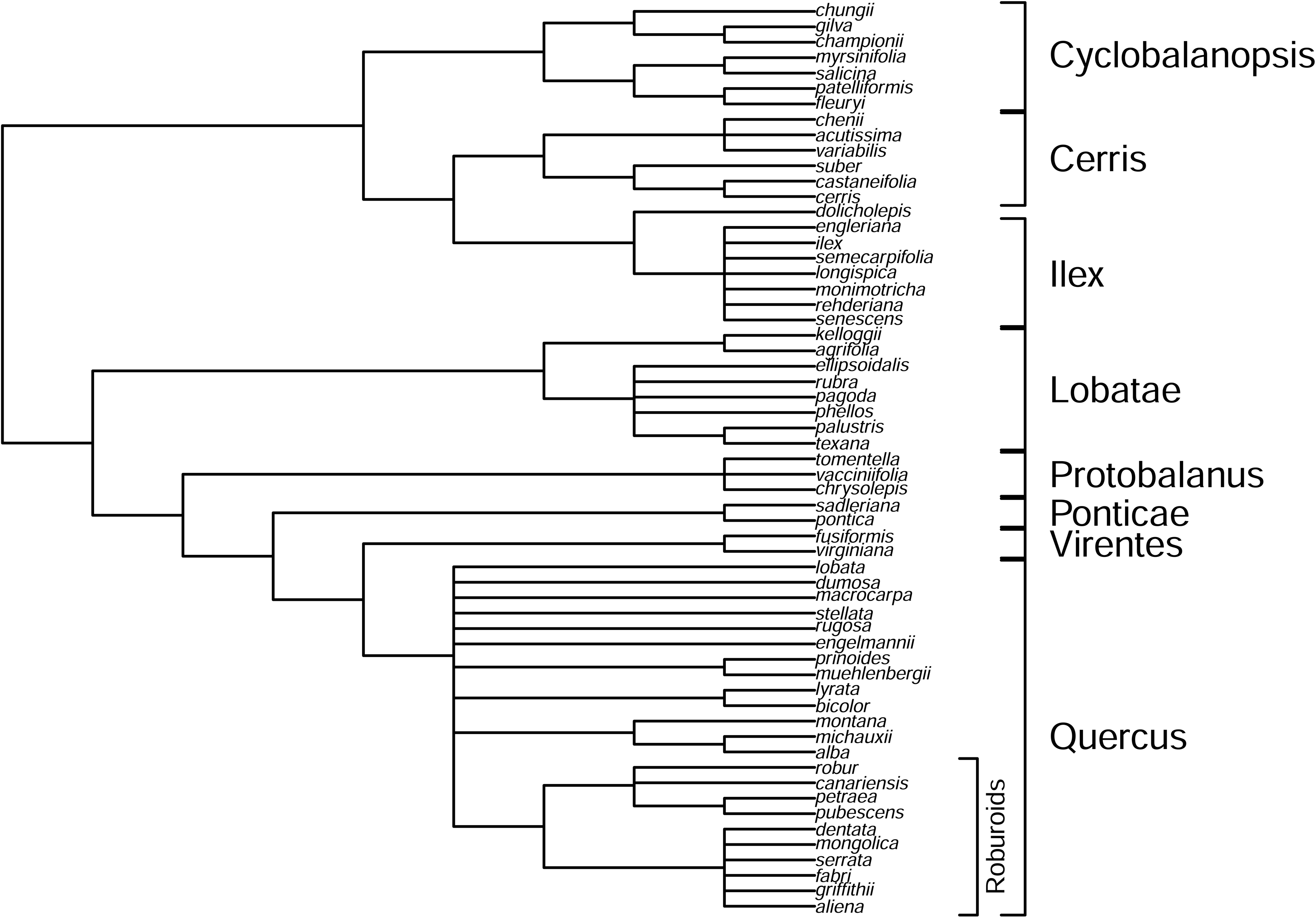
Strict consensus of all ML trees across all analyses and datasets in this study, excluding *de novo* clustering of empirical RADseq data. Trees are pruned to only taxa common across analyses and datasets. *De novo* clustering tree is excluded because it is the only analysis in which sect. *Quercus* is paraphyletic (see text). Branch lengths are arbitrary; tip labels are all *Quercus* epithets. Clade labels indicate the eight *Quercus* sections, whose positions are resolved consistently across all datasets and analyses; and the Roburoid clade of white oaks, which comprises the European white oaks (*Q. robur* through *Q. pubescens* in this figure) and the East Asian white oaks (*Q. dentata* through *Q. aliena* in this figure).

#### Gene flow inferences: *f*-branch analyses

Analysis of phylogenetic discordance using the f-branch metric (*f*_b_) based on the *Q. alba* reference shows that introgression is not rampant across the oak phylogeny, but scattered, with a few high-introgression areas of the tree (Fig. 5). We summarize introgression inferences from the *f*-branch metric (*f*_b_) using the maximum *f*_b_ by row, given the propensity of *f*_b_ plots to show rows of high values due to shared introgressed alleles among close relatives (Malinsky *et al*., 2021). The two taxa with highest *f*_b_ values are *Q. engleriana* (sect. *Ilex*) with the other East Asian members of sect. *Ilex* (max *f*_b_ = 0.173), and its reciprocal comparison, the branch subtending the east Asian members of sect. *Ilex* with *Q. engleriana* (max *f*_b_ = 0.110); and *Q. pontica* (sect. *Ponticae*) with the Eurasian white oaks (“Roburoids” of sect. *Quercus*; max *f*_b_ = 0.094), with its reciprocal comparison, the ancestor of all white oaks with *Q. pontica* (max *f*_b_ = 0.070). Two additional high *f*_b_ values bear noting: *Q. engelmannii*, a white oak endemic to southern California that originates from the Mexican white oak clade (Hipp et al., 2018), shows significant evidence of introgression with *Q. dumosa* (a relatively low but significant *f*_b_ = 0.047); and *Quercus macrocarpa* shows significant allele sharing with *Q. lobata* (max *f*_b_ = 0.077), and *Q. lobata* with *Q. dumosa* (max *f*_b_ = 0.075).

**Figure 5.**
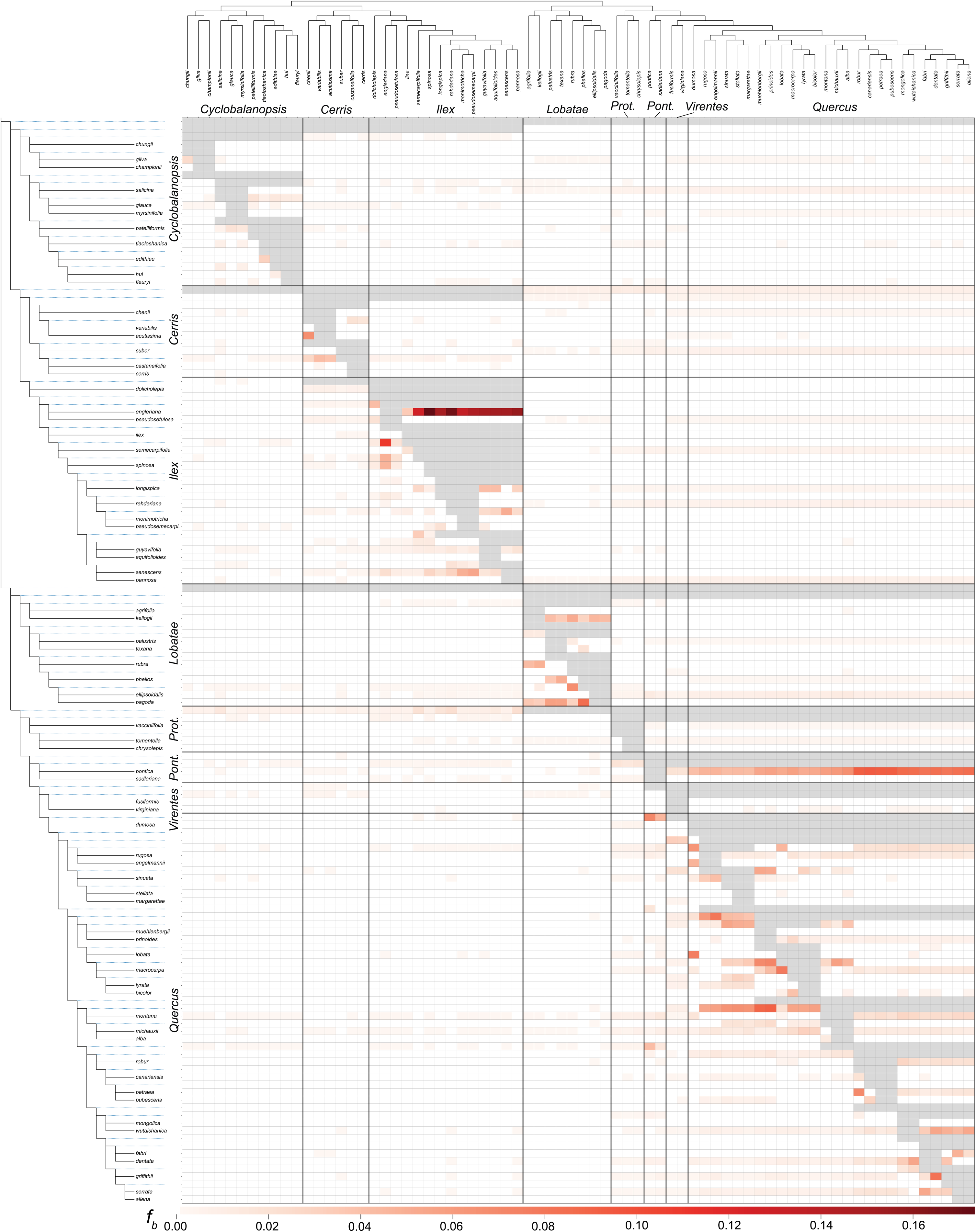
Heatmap of f-branch (*f*_b_) metric summarizing phylogenetic discordance in the whole genome dataset based on the tree estimated from *Q. alba*-mapped SNPs. Cells indicate excess allele-sharing between tips (columns of the matrix) and either nodes or tips of the phylogeny (rows of the matrix), relative to the sister species to each node or tip (in the phylogeny on the left edge of the figure). All colors other than white are considered significant (*p* < 0.01; details in text); gray cells are pairwise comparisons that cannot be tested due to the tree topology used. Because allele sharing is relative to a sister species based on phylogenetic invariants, significant allele-sharing is treated as evidence of introgression, though significant values may also be influenced by among-lineage rate variation. Section names are provided along the margins.

Summarized over all tip-branch comparisons in which a branch falls within a single section—that is, excluding *f*_b_ values that compare a tip with a branch whose descendents are from more than one section—average intersectional *f*_b_ values (*f*_b_ = 0.00070 +/- 0.00005 [S.E.M.], *n* = 7,644) are significantly lower than intrasectional *f*_b_ values, which range from 0.0017 +/- 0.0003 (*n* = 231) in sect. *Cyclobalanopsis* to 0.0146 +/- 0.0020 (*n* = 120) in sect. *Lobatae* and 0.0143 +/- 0.0018 (*n* = 378) in sect. *Ilex* (Table 1). Conducting the *f*-branch analysis with sections as tips similarly demonstrates that all pairwise *f*_b_ values are quite low, except for one: section *Ponticae* shares significantly more alleles with section *Quercus* than with the sister section *Virentes* (Fig. 6).

**Figure 6.**
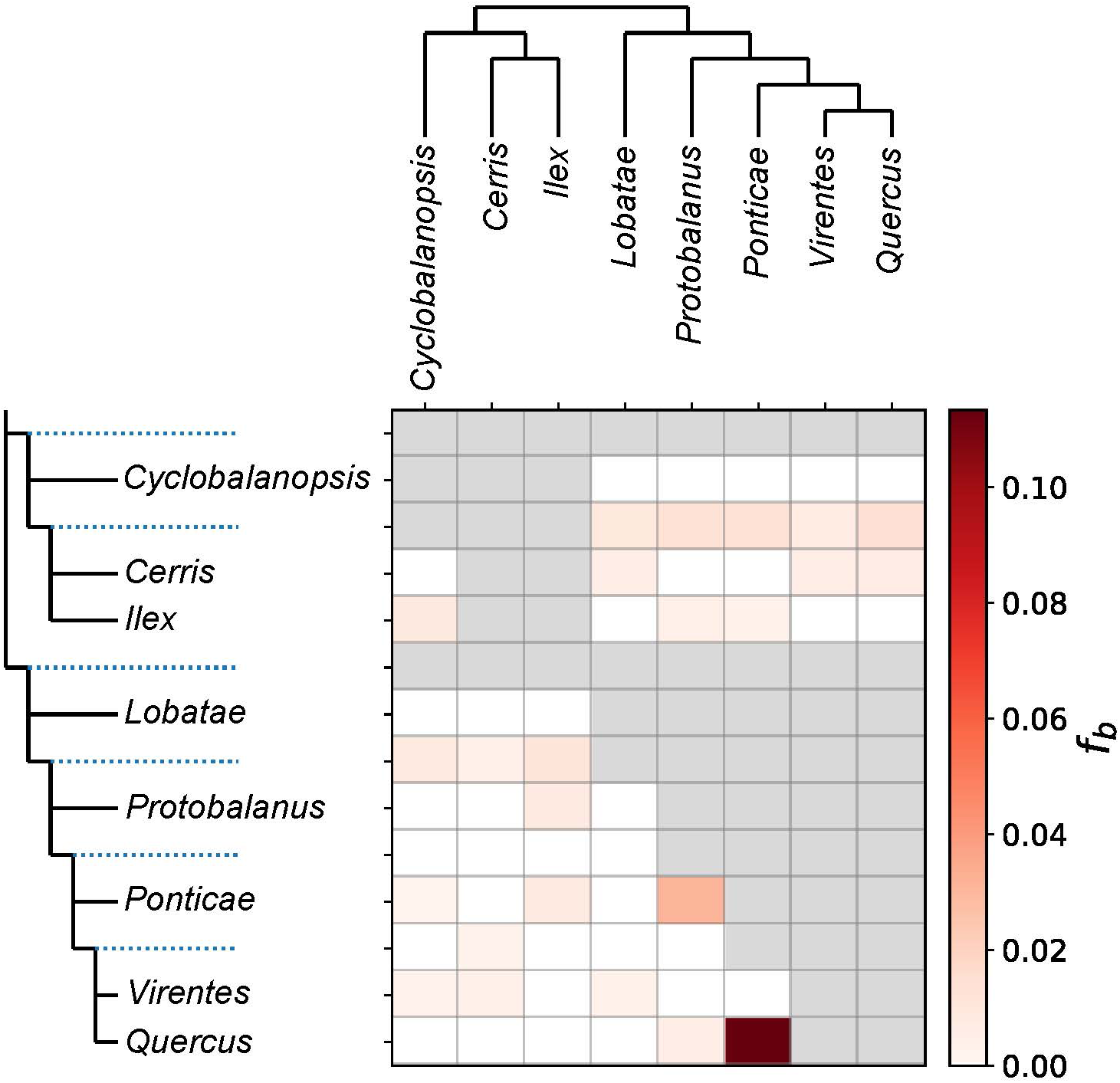
Heatmap of f-branch (*f*_b_) metric summarizing among-section phylogenetic discordance at the section level. This analysis is based on the same tree and dataset used for *f*_b_ statistics of species (Fig. 5). In this analysis, however, species are treated as exemplars of the section, so that the *f*_b_ values estimate gene flow between sections, averaged over species.

**Table 1.**
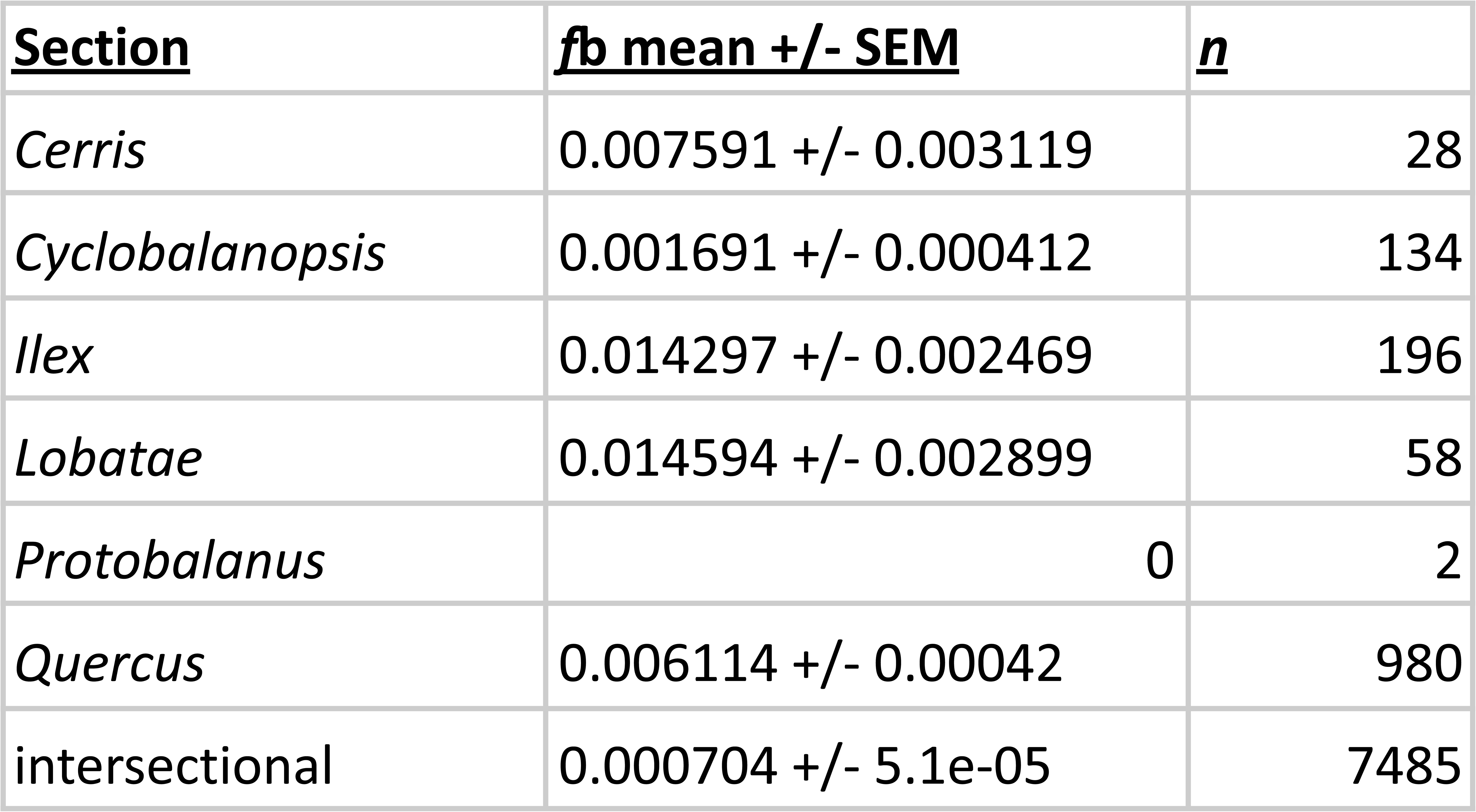
F-branch summaries by section. All pairwise *f*_b_ values are averaged for each branch of the phylogeny as described in the text, treating non-significant values (*p* > 0.01) as 0. Comparisons in which the branch subtended a mixture of sections were excluded (*n* = 153), leaving a total of 8,883 comparisons. “Intersection” denotes comparisons in which the branch compared subtends individual(s) of one section, and the tip to which it is being compared is from a different section. Sections *Virentes* and *Protobalanus* were represented by two tips only, so no *f*_b_ values could be calculated for them.

A relative-rate test based on the *Q. alba* reference reveals substantial heterogeneity in the molecular rates of evolution across the oak phylogeny. The largest rate disparity is concentrated in sect. *Ilex* and sect. *Virentes* (Fig. S4). The two species with the highest mean rate difference against all other species are *Q. pannosa* (mean rate difference = 0.410) and *Q. rehderiana* (mean = 0.405). *Quercus semecarpifolia* (sect. *Ilex*; mean = 0.357), *Q. virginiana* (sect. *Virentes*; mean = 0.378), and *Q. fusiformis* (sect. *Virentes;* mean = 0.337) similarly show markedly elevated rates relative to the rest of the genus. By contrast, the slowest evolving lineages are the eastern North American red oaks (sect. *Lobatae*: *Q. rubra, Q. phellos*, *Q. palustris*, and *Q. pagoda*; mean = 0.20), the california red oak *Q. kelloggii* (mean = 0.13), the Eurasian sect. *Cerris* (*Q. cerris*, *Q. variabilis*; mean = 0.14), *Q. alba* (sect. *Quercus*; mean = 0.141), and *Q. pontica* (sect. *Ponticae*; mean = 0.145).

At the section level, pairwise mean rate differences are dominated by comparisons with section *Virentes* (Fig. S4). The highest mean rate differences are between *Protobalanus*-*Virentes* (mean = 0.479), *Ilex-Virentes* (0.458), *Cyclobalanopsis-Virentes* (0.377), *Ponticae-Virentes* (0.362), and *Quercus-Virentes* (0.346). Sect. *Ilex* also shows elevated rate differences against sect. *Cerris* (0.332), sect. *Quercus* (0.294), sect. *Ponticae* (0.267), and sect. *Lobatae* (0.266). The most rate-similar sections are *Cyclobalanopsis-Protobalanus* (0.057), *Cerris-Quercus* (0.057), *Cerris-Ponticae* (0.063), *Cerris-Lobatae* (0.064), and *Lobatae-Ponticae* (0.064), indicating broadly comparable evolutionary rates among these clades. Within sections, intrasectional rate variation is highest in sect. *Ilex* (mean = 0.249 ± 0.017 (SE)], n = 91 comparisons) and sect. *Lobatae* (0.203 ± 0.029, n = 28), and lowest in sect. *Cyclobalanopsis* (0.087 ± 0.011, n = 55) and sect. *Quercus* (0.100 ± 0.004, n = 325). Across all between-tip comparisons spanning two different sections, the overall mean intersectional rate difference is 0.196 ± 0.003 (n = 2,037).

### Comparing trees inferred from different datasets

Ordination of all trees pruned down to the 59 common taxa recovers three major phylogenetic ordination groups: (1) one large group comprising the bootstraps and ML trees for the simulated RADseq (‘simRAD’) and (2) one comprising the empirical RADseq (‘empiricalRAD’ and ‘*de novo*’) analyses; these two phylogenetic groups vary parallel to each other. (3) A third small ordination group comprises the ML trees and bootstraps of the genome resequencing analyses, near one end of the simRAD ordination group (Fig. 7). The genome resequencing bootstrap trees are all so similar—many identical to one or the other ML tree—that they are clustered too tightly to be discerned in the ordination (Fig. 7). The previously published tree of Hipp et al. (2020) falls close to the empirical RADseq bootstraps (“OakPhylo2020” in Fig. 7), presumably because this analysis was constrained in light of previous analyses demonstrating the effects of *Ponticae* – *Quercus* introgression as well as the observation that in the global analysis, *Quercus* was paraphyletic with respect to *Virentes*. The most significant differences in clades distinguishing the genome resequencing from the simRAD trees are the monophyly of the California white oaks sampled (0% in genome resequencing, 36.2% in simRAD; Table S5, S6) and the position of the Roburoid white oaks (sister to subsection *Albae* in 100% of genome resequencing bootstraps, 37.2% of simRAD bootstraps; Table S6).

**Figure 7.**
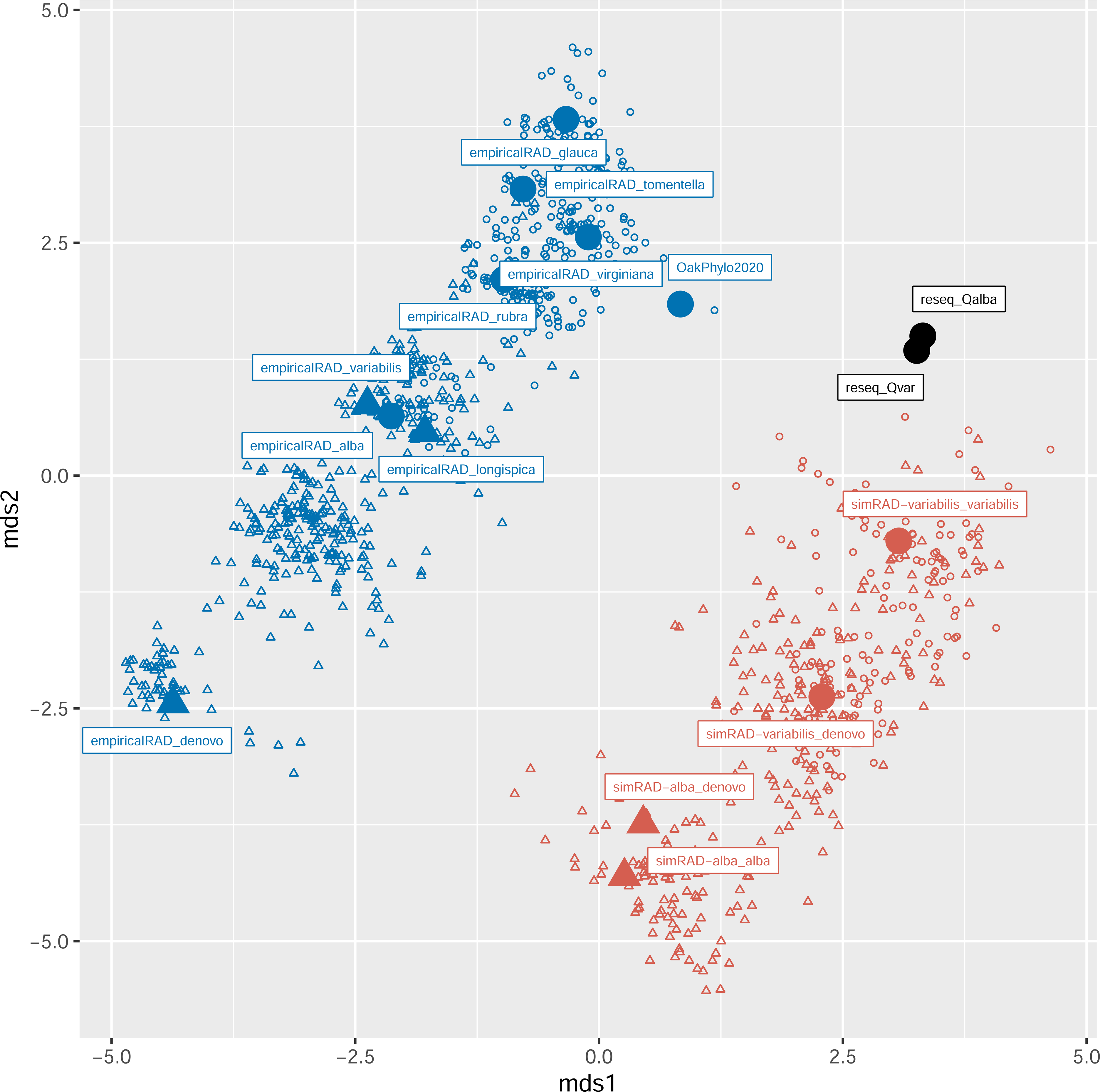
Ordination of phylogenies using metric multidimensional scaling on pairwise entropy distances. Ordination was conducted at *K* = 2 dimensions. **Symbols:** Larger, filled symbols indicate ML trees; smaller, open symbols represent bootstrap replicates from the same dataset as the ML trees. Circles indicate trees in which the Roburoid white oaks fall sister to subsection *Albae*; triangles indicate trees in which the Roburoid white oaks fall sister to all remaining white oaks. **Abbreviations:** reseq = SNP data based on whole genome resequencing dataset; simRAD = simulated RADseq, based on the pseudo-reference sequences dataset; empiricalRAD = empirical RADseq mapped back to a genome; denovo = RADseq clustered *de novo*, without a reference genome; OakPhylo2020 = phylogeny of Hipp et al. 2020. In all cases, the specific epithet of the reference species used for SNP-calling and locus-assembly is indicated after the underscore.

Three broad patterns stand out in the tree comparisons. First, the simRAD trees are biased with respect to the trees generated from the full genome resequencing datasets, despite the fact that the simRAD data are generated from the pseudo-reference genomes created using the resequencing data. This is evidenced by the fact that the cloud of bootstrap trees from the simulated RADseq data do not include the trees generated from the full genome resequencing data. The reason for this is not immediately obvious: while the genome resequencing bootstrap trees exhibit lower variance (less spread in the ordination), the strict consensus of all the simRAD bootstraps (34 resolved nodes) is entirely compatible with the strict consensus of all genome resequencing bootstraps (69 resolved nodes in the genome resequencing bootstrap strict consensus; 34 resolved nodes in the combination of all simRAD and genome resequencing bootstraps). No single relationship stands out in the genome resequencing vs simRAD trees, yet the combination of resolved nodes is unique: the Robinson-Foulds distance between the genome resequencing trees and every non-reseq bootstrap is > 0.

Second, the genome resequencing and simRAD trees (Fig. 3, S2, S3) resolve the Mexican *Quercus* (including *Q. engelmannii*) as forming a clade with the Stellatae, while the empirical RAD trees (Fig. 2, S1) mostly do not (Table S5). The two sampled oaks of Mexican origin (*Q. engelmannii* and *Q. rugosa*) fall sister to each other in 100% of the combined genome resequencing and simRAD bootstrap trees, but only 2.9% of the empirical RADseq bootstrap trees (Table S6). Additionally, 91.7% of the combined genome resequencing and simRAD bootstraps resolve the Mexican white oaks and subsection *Stellatae* as a clade, while only 19.1% of the empirical RADseq bootstraps do (Table S6).

Finally, the bootstrap sets for the simulated RAD and the empirical RAD exhibit a comparable spread (area covered in the 2-dimensional ordination space), and the longest axis of variation in both is explained by whether the Roburoids are sister to subsection *Albae* (circles in Fig. 7) or sister to the remainder of sect. *Quercus* (triangles in Fig. 7). The empirical RADseq ML trees sort into two overlapping ordination clusters: one in which the Roburoids fall sister to the remainder of the white oaks (mapped to references *Q. glauca*, *Q. rubra*, and *Q. virginiana*), and one in which the Roburoids fall sister to subsection *Albae*. The variation within each bootstrap set is quite high: the average bootstrap support for the Roburoid-*Albae* clade in the *Q. alba*, *Q. glauca*, *Q. rubra*, *Q. tomentella*, and *Q. virginiana* datasets, whose ML trees support this relationship, is only 67%. Support is as low as 36% in the dataset mapped to *Q. tomentella* (Fig. 2; Table S6). As in oak populations, the within-group variation—in this case, phylogenetic variation within bootstrap sets—is very high in comparison to among-group variation—the differences between bootstrap sets derived from a single dataset.

One important caveat in these comparisons is that the empirical RADseq datasets comprised 322 taxa and were pruned down post-phylogenetic inference. This may introduce a systematic difference between the empirical and simulated RADseq trees that is not explored in this chapter.

## Discussion

This chapter spotlights three sources of conflict among oak phylogenetic inferences: library preparation / data type (RADseq vs whole genome resequencing); the reference genome used for variant-calling; and hybridization / introgression. The latter two effects explain much of the uncertainty in RADseq phylogenies as assessed by bootstrapping (Fig. 7). The study also demonstrates that there is broad congruence in the topology of the oak phylogeny based on all genome datasets we assess (Fig. 4). This gives us good reason to be confident that our growing understanding of oak phylogenetic history rests on a strong foundation.

### Data types: RADseq vs whole genome resequencing

There are two major differences in phylogenies estimated from whole-genome resequencing data and simulated RADseq (simRAD). First, despite the fact that simRAD data are derived from the resequencing data, the simRAD bootstrap set does not overlap with the resequencing topologies (Fig. 7). In other words, the tree inferred from resequencing data contains a combination of nodes that is not found in any of the simRAD bootstraps. This result is surprising: we had expected the whole-genome resequencing data to result in a tree with low variance—i.e., tightly clustered bootstrap trees, as we do find—but more or less central to the simulated RADseq (simRAD) bootstraps. While every bootstrap tree in the resequencing dataset supports Roburoids sister to subsection *Albae*, *Q. macrocarpa* sister to *Q. bicolor*, and *Q. lobata* in a clade with *Q. macrocarpa* and *Q. bicolor*, the simRAD data support these relationships in only 37.3%, 41.8%, and 34.8% of bootstrap trees, respectively (Table S6). Moreover, only 1.0% of simRAD trees recover the same combination of key white oak clades found in every one of the whole-genome resequencing bootstraps (Table S5). Our results consequently suggest that the RADseq and whole genome sequencing datasets produce trees that differ systematically, not just in precision.

Does whole genome resequencing “win” over RADseq? We are inclined to say yes, but with a caveat. For sheer information content, whole genome resequencing data is the winner, hands down: on average, the resequencing datasets had 63.0× as many site patterns as the empirical RADseq data (Table S4b; the simRAD datasets were comparable to the empirical RADseq datasets, at 3.0× as many site patterns on average). However, while whole genome resequencing data enable gene-by-gene exploration and modeling of phylogenomic discordance, many analyses, even of whole-genome data, are still performed on concatenated SNP matrices, as we have done in this chapter. For such analyses, we consider the higher phylogenetic variance found in the RADseq datasets as a potential benefit, as that variance can pinpoint areas of gene tree discordance.

The placement of the California white oak *Q. lobata* is an interesting cast-study in how RADseq and whole-genome resequencing data provide different phylogenetic insights. This species falls in a clade sister to or near the eastern North American *Q. macrocarpa* in all whole-genome analyses with seemingly perfect support: whole-genome resequencing data soundly reject monophyly of the true California white oaks based on the two species in our datasets (*Q. lobata* and *Q. dumosa*; 0% bootstrap support). Yet the simRAD dataset suggests ambiguity (36.3% support for *Q. lobata* and *Q. dumosa* monophyly as the only two California white oak clade members sampled in this study). Our previous analyses of empirical RADseq data (McVay *et al*., 2017a; Hipp *et al*., 2018) recovered *Q. lobata* as sister to the remaining California white oaks—as our *de novo* RADseq dataset does in this study—and demonstrated a high degree of allele-sharing between *Q. gambelii* (not represented in this study) and both *Q. lobata* and *Q. macrocarpa*, causing instability in the white oaks. The species-level f-branch analysis presented in this chapter (Fig. 5) shows gene flow between *Q. lobata* and both *Q. dumosa* and *Q. macrocarpa*, suggesting that the high confidence for placement of *Q. macrocarpa* in the whole genome tree belies a mosaic genomic history resulting from ancient hybridization between taxa that are very distantly allopatric today. That mosaic history would be more easily dissected using a granular, gene-by-gene approach on whole genome data (Crowl *et al*., 2020; e.g., Thawornwattana *et al*., 2023; Shen *et al*., 2025).

The uncertainty of RADseq is, of course, not what one hopes for in a dataset, but it can help point to introgression histories that bear further investigation.

### Choice of reference genomes

Recent work demonstrates that use of any particular reference genome in variant-calling affects the SNP composition of the resultant dataset and inferences based on that dataset (Thorburn *et al*., 2023; Mohn *et al*., 2026). In our study, mapping whole-genome data to different reference genomes that are separated by the deepest split in the oak phylogeny—*Q. alba* is in subgenus *Quercus*, *Q. variabilis* in subg. *Cerris*—has negligible effect on phylogenetic inference (Fig. 3). In the empirical RADseq data, trees do not cluster more closely based on the relatedness of the reference used as assessed either by ordination of bootstraps (Fig. 7) or visual inspection of the trees (Fig. 2). This may be due in part to our use of BWA, which has relatively low stringency to the reference (Mohn *et al.,* 2026).

It is striking that *de novo* clustering of the empirical RADseq data recovers an average of 28.4% more loci than the empirical RADseq datasets with reference mapping (Table S4a) while in the simulated RADseq dataset, *de novo* clustering and reference-guided clustering recover approximately the same number of loci (Table S4a). It is unclear why the concatenation of adjacent loci in the reference-guided datasets does not result in a large decrease in loci in the simRAD datasets; whether sequence coverage may explain this bears exploring in future analyses. Also worth exploring is whether any of the additional loci in the *de novo* clustering of the empirical RADseq data are due to non-nuclear DNA. The chloroplast is likely to be a minor contributor: in an early publication from a portion of this dataset, we verified that there are only 13–14 *Pst*I restriction sites in the oak chloroplasts investigated, and only 39 of 109,992 loci from *de novo* clustering mapped to known oak chloroplasts (McVay *et al*., 2017b). Whether these extra loci may include contaminants (such as fungal or bacterial) filtered out by the reference, or whether they are real loci that are missing from the *Q. alba* reference (which would also be filtered out in the simulation process) remains to be seen in future analyses.

### Effects of introgression on topological variation

Without quantifying the relative effects of ILS, gene tree estimation error, and introgression in explaining the variance with bootstrap datasets or among analyses (e.g., Liu *et al*., 2025; Shen *et al*., 2025), our analyses demonstrate that much of the variation among phylogenetic inferences in the genus can be explained by hybridization. Among datasets, the simRAD and whole genome datasets place *Q. engelmannii* sister to the other Mexican white oak sampled in all datasets (*Q. rugosa*) (Fig. 3, S3), whereas the empirical RADseq place it among the California white oaks (Fig. 2). This major point of difference between the datasets is shaped by ancient introgression. In the whole genome resequencing data, *Q. engelmannii* shows an excess of allele sharing with the Californian *Q. dumosa* (Fig. 5). *Quercus englemannii* is morphologically extremely similar to the Mexican *Q. oblongifolia* (Nixon, 1997), to which it has fallen sister in previous RADseq analyses (Hipp *et al*., 2020; Althaus *et al*., 2026) of this same dataset, using *de novo* clustering. Population-level analyses have demonstrated that *Q. engelmannii* exhibits significant introgression from the Californian scrub white oak *Q. berberidifolia* (Kim *et al*., 2018). Even introgression from *Q. berberidifolia* into *Q. engelmannii* is widespread and not limited to sites where the two species are sympatric, *Q. engelmannii* individuals vary in the amounts of introgression they exhibit (O’Donnell *et al*., 2021). In previous analyses, we have found *Q. engelmannii* shifting between the California white oak and Mexico white oak clades depending on the individual sampled as well as clustering parameters. Analyses in this chapter support the inference that this instability is due to the ancient introgression detected in other work (cited above).

Variation within each dataset is also strongly influenced by a single prominent difference: placement of the Eurasian (Roburoid) white oaks (Fig. S5). Phylogenetic affiliation of the Roburoids was studied in detail in previous work (McVay *et al*., 2017b; Crowl *et al*., 2020; Zhou *et al*., 2022), which demonstrated that the Roburoids are sister to a trio of eastern North American white oak species (subsection *Albae*), but phylogenetically unstable due to introgression. By one estimate, Roburoids may contain ca. 11% of their genome from ancient gene flow with *Q. pontica* of the Caucasus (Crowl *et al*., 2020). Variation in the placement of this clade—either sister to the eastern North American *Albae* or sister to the remainder of the section—correlates very strongly with the long axis of variation in each RADseq ordination cluster (Fig. 7, S5).

Thus introgression is responsible for both among-dataset and within-dataset variation in our study. Our work suggests that rate variation among clades, particularly involving section *Virentes*, may bias Type I error rates in some of the f-branch statistics calculated (Frankel & Ané, 2023; Pang *et al*., 2025). We suspect our results are minimally impacted, both because most introgression events we observe within our study are within sections, and the relative rate tests suggest no significant difference in substitution rates between sections *Quercus* and *Ponticae* (the major area of intersectional variation within the study). To the contrary, *Q. pontica* has one of the slower substitution rates, which reduces the risk of artifactually detecting shared alleles between it and non-sister lineages (e.g., the Eurasian white oaks, where it shows a high surplus of shared alleles).

## Conclusions: where do we stand on the oak phylogeny?

Our results suggest that the backbone tree topologies used in previous analyses (e.g., Hipp et al. 2020, Althaus et al. 2026) are stable and well supported by both RADseq and whole-genome resequencing. Mohn et al. (2026) has shown that reference genome choice affects heterozygosity, branch lengths, and tree topology with stringent mapping criteria and at small scales within the eastern North American white oaks. Our results suggest that at the genus scale with more relaxed mapping criteria, the tree topology biases are minor relative to the phylogenetic signal in the data. Additionally, the constraints used in the prior studies (“OakPhylo2020 in Fig. 7) are largely supported by whole-genome resequencing, which recovers most of the same major nodes (Table S5).

*Quercus* section *Quercus*—the white oak clade—stands out as the most unstable part of the oak phylogeny. This is a long-standing problem. In Manos et al. (1999), the Mexican white oaks did not form a clade. In Denk & Grimm (2010), Roburoid white oaks do not form a clade. In Huber et al. (2014), the eastern North American, western North American, and Mexican white oaks are intermixed, and *Q. pontica* is embedded among the Roburoid white oaks (with 6% bootstrap support for the clade). The major points of phylogenetic discordance within the section are placement of the Roburoid clade (i.e. the Eurasian white oaks), documented in RADseq (McVay et al. 2017; Hipp et al. 2018, 2020), sequence capture (Crowl et al.2020) and resequencing data (Zhou et al. 2022; this study); and placement of *Q. lobata* (investigated in McVay *et al*., 2017a), which moves between the eastern North American and western North American white oak clades depending on the analysis. Both of these issues bear additional study with closer interrogation of whole-genome data.

For phylogenetic inference, RADseq data will likely be eclipsed by whole genome sequencing data, particularly as the costs of genome resequencing and computation and storage costs continue to drop. Estimating tree phylogenies in the face of ILS, gene tree estimation error, and introgression requires whole genome sequencing data or large numbers of loci. Even so, RADseq has valuable roles in the near future for fine-scale phylogenetics, at the boundaries between population differentiation and speciation, and in studies of species-boundaries and introgression. Additionally, combinability with existing data, affordability, and computational tractability continue to make RADseq useful, and the analyses in this chapter give us good reason to be confident in its utility.

## Acknowledgments

We are very grateful to Ilga Porth and Yousri El-Kassaby for inviting us to submit this chapter, and for their work editing this exciting volume. This chapter would not be possible without the support of collaborators on the many previous RADseq phylogenies in which portions of these data were used. It would also not be possible without support of the Morton Arboretum Herbarium, particularly Lindsey Worcester and the herbarium volunteers, in archiving specimens and making them available online. Special thanks to Matthew Hahn for many discussions that helped in developing the workflow for processing the resequencing data. The authors gratefully acknowledge the use of cyberinfrastructure managed by the Research Services Department of UITS at Indiana University. Through the use of these computing resources, this research was supported in part by Lilly Endowment, Inc., through its support for the Indiana University Pervasive Technology Institute. The manuscript benefited from feedback from staff, students, and affiliates of the Herbarium and Systematics Lab at The Morton Arboretum. ALH was supported NSF Award #1146488 and a Fulbright Fellowship funded by the Franco-American Commission; RM by NSF Award #2129281; and DAL by NSF IOS-2109716.

## Supplement: captions

Table S1. Taxon sampling for RADseq data.

Table S2. Taxon sampling and sequencing statistics for whole genome resequencing data.

Table S3. Simulated RADseq empirical clustering stats summary.

Table S4a. RADseq loci in assembled in ipyrad for each dataset.

Table S4b. Phylogenetic site patterns per dataset.

Table S5. Clade presence / absence for all maximum likelihood (ML) trees.

Table S6. Percent of bootstrap trees resolving each of the major clades discussed in text.

Figure S1. RADseq empirical phylogenies by reference genome. These correspond with the section-level phylogenies (Fig 2, main text) but include full taxon sampling. Reference genomes used: (a) *de novo* clustering [no reference], (b) *Quercus alba*, (c) *Q. glauca*, (d) *Q. longispica*, (e) *Q. rubra*, (f) *Q. tomentella*, (g) *Q. variabilis*, (h) *Q. virginiana*.

Figure S2. Whole genome resequencing (SNP) phylogenies, all taxa (ingroup and outgroup). Reference genomes used for variant calling: (a) *Quercus alba*, (b) *Q. variabilis*.

Figure S3. Simulated RADseq phylogenies by reference genome. Reference genomes used: (a) *de novo* clustering [no reference], (b) *Quercus alba*, (c) *Q. glauca*, (d) *Q. longispica*, (e) *Q. rubra*, (f) *Q. tomentella*, (g) *Q. variabilis*, (h) *Q. virginiana*.

Figure S4. Heatmap of f-branch (*f*_b_) metric summarizing among-section phylogenetic discordance at the section level, with associated rate-difference test.

Figure S5. Clade loadings on ordination (cf. Fig. 7).

## Notes

https://zenodo.org/records/20192671

https://github.com/andrew-hipp/oakphylo2026

### Competing Interest Statement

The authors have declared no competing interest.

