## Supplementary figures and images for "Resolving the oak tree of life: comparing RADseq and whole genome resequencing methods for oak phylogenetics"

### Fig. S1a

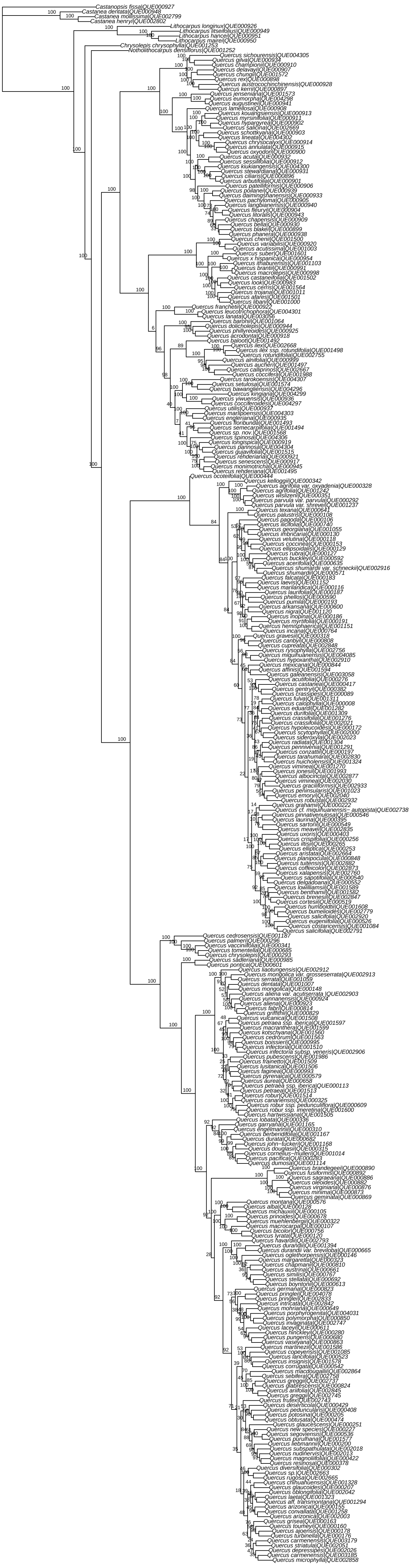

### Fig. S1b

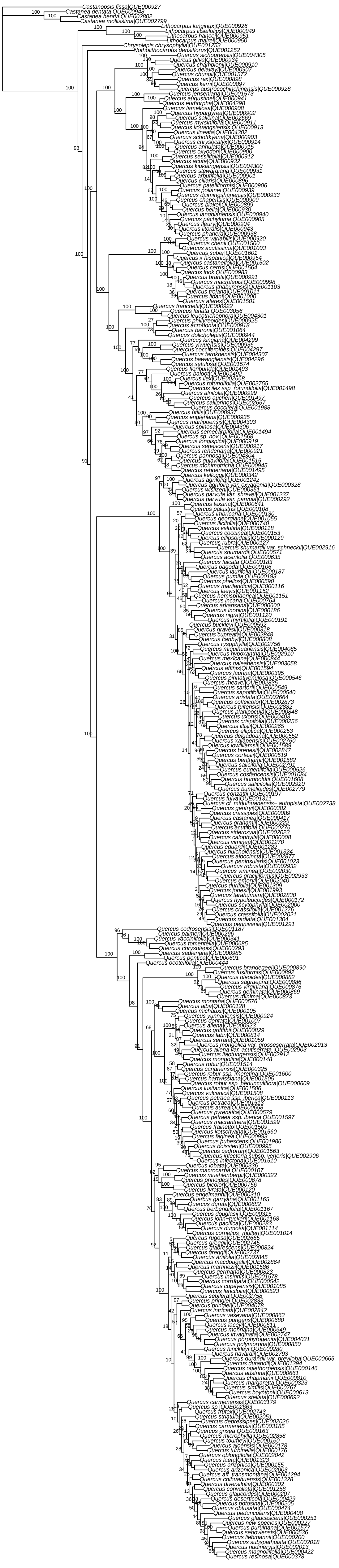

### Fig. S1c

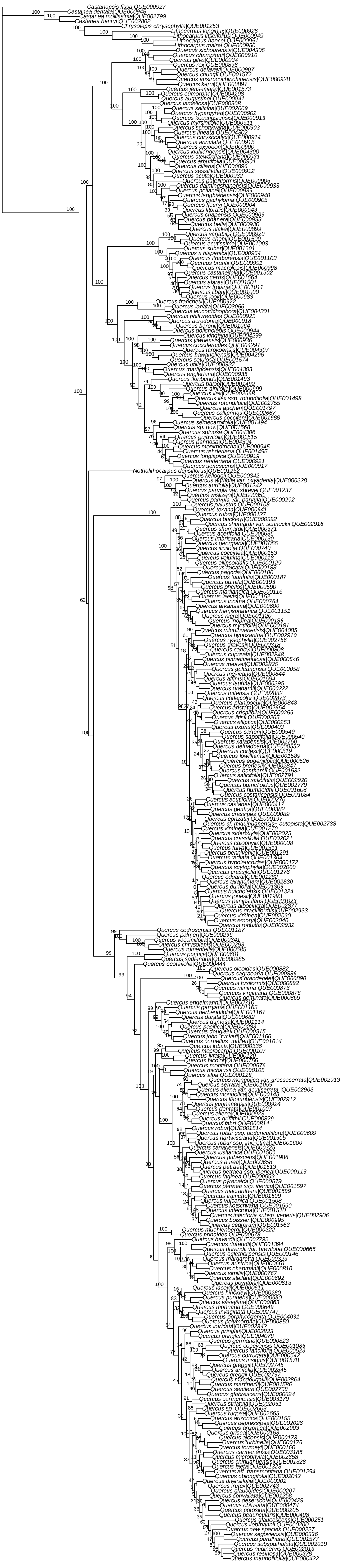

### Fig. S1d

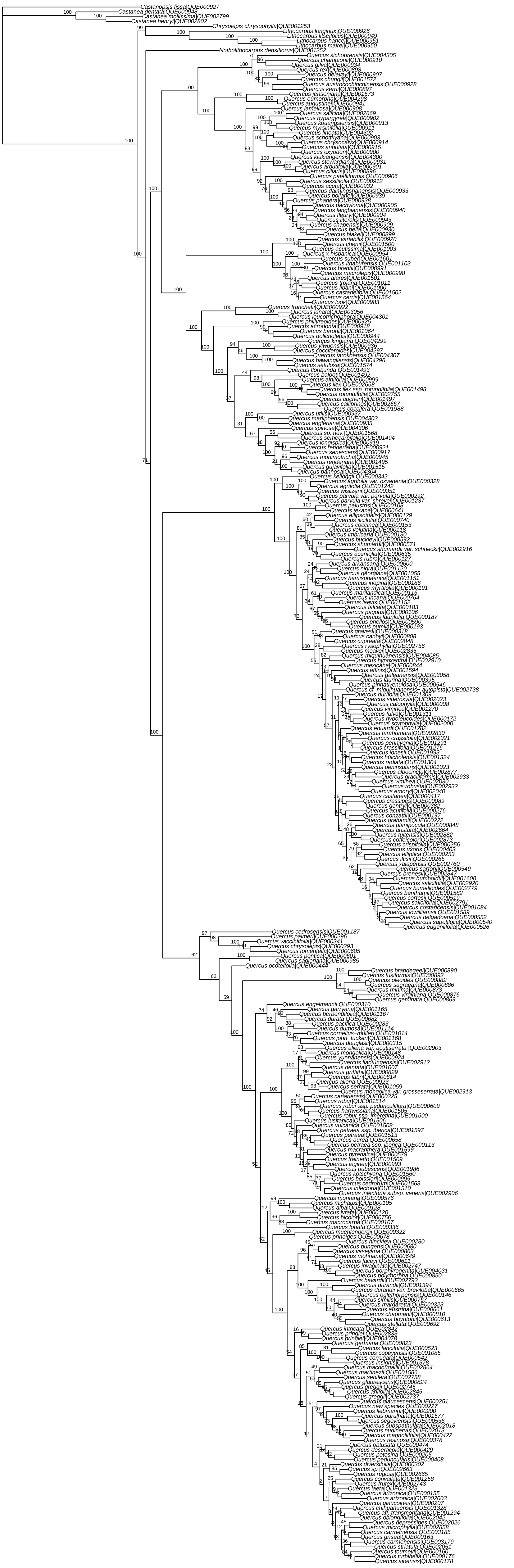

### Fig. S1e

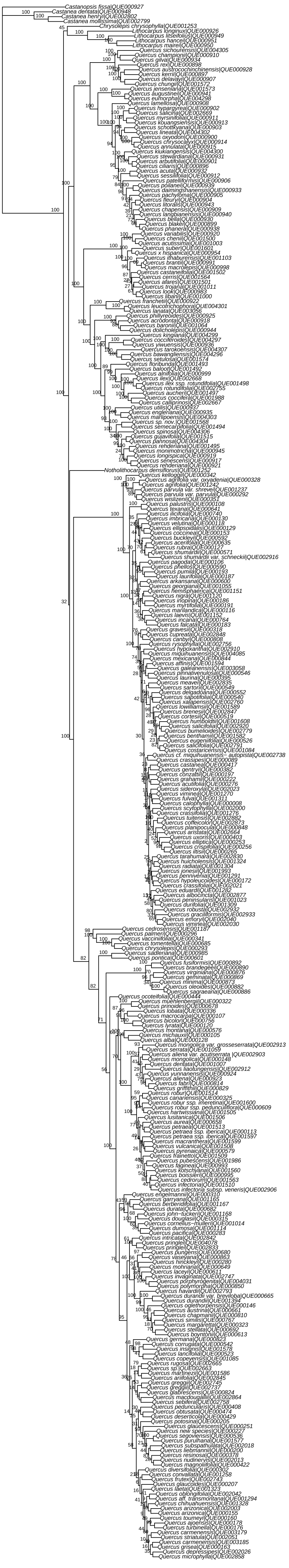

### Fig. S1f

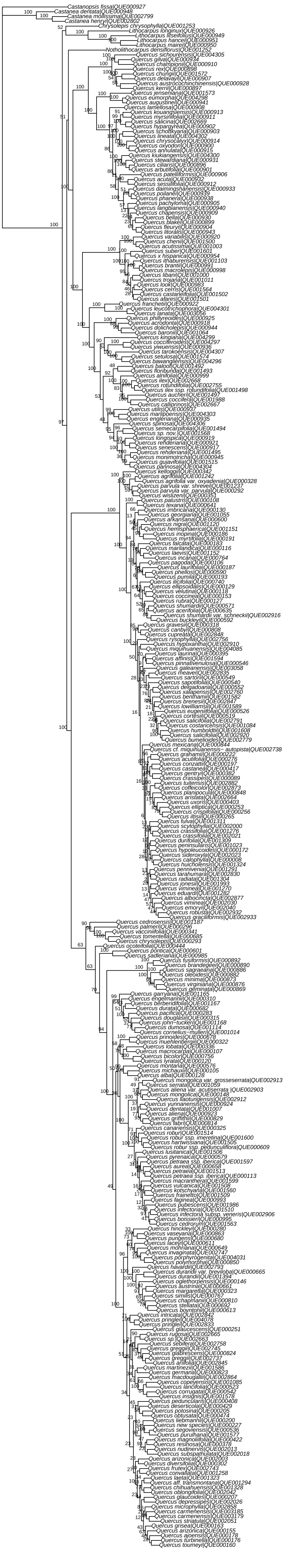

### Fig. S1g

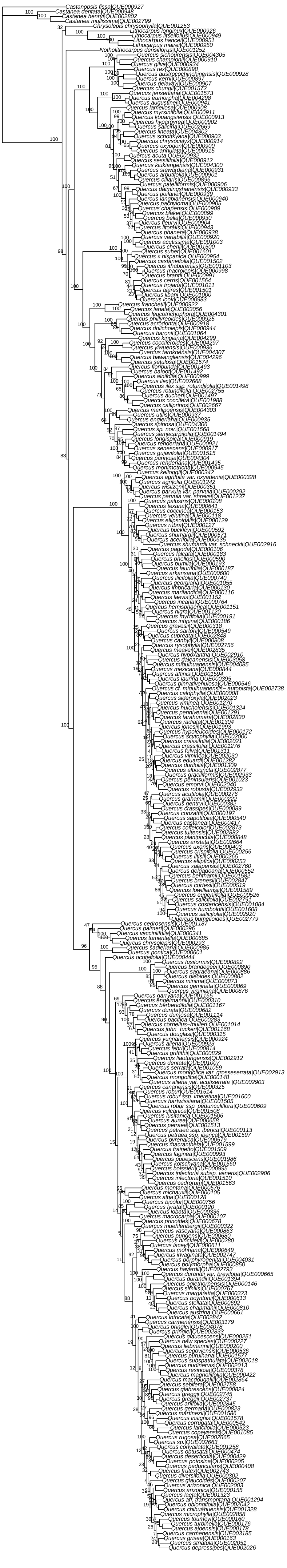

### Fig. S1h

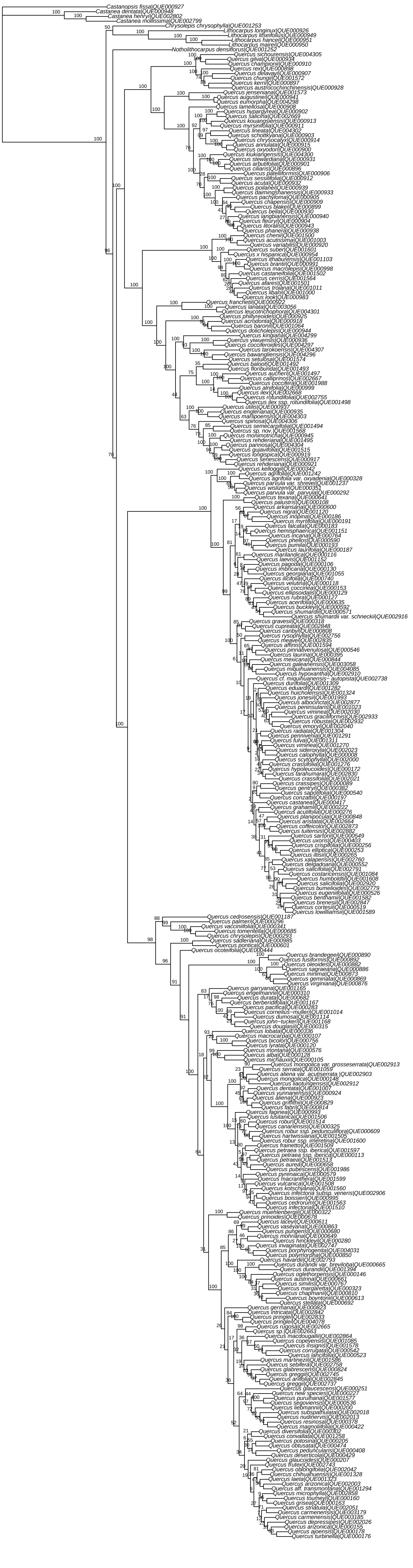

### Fig. S2a

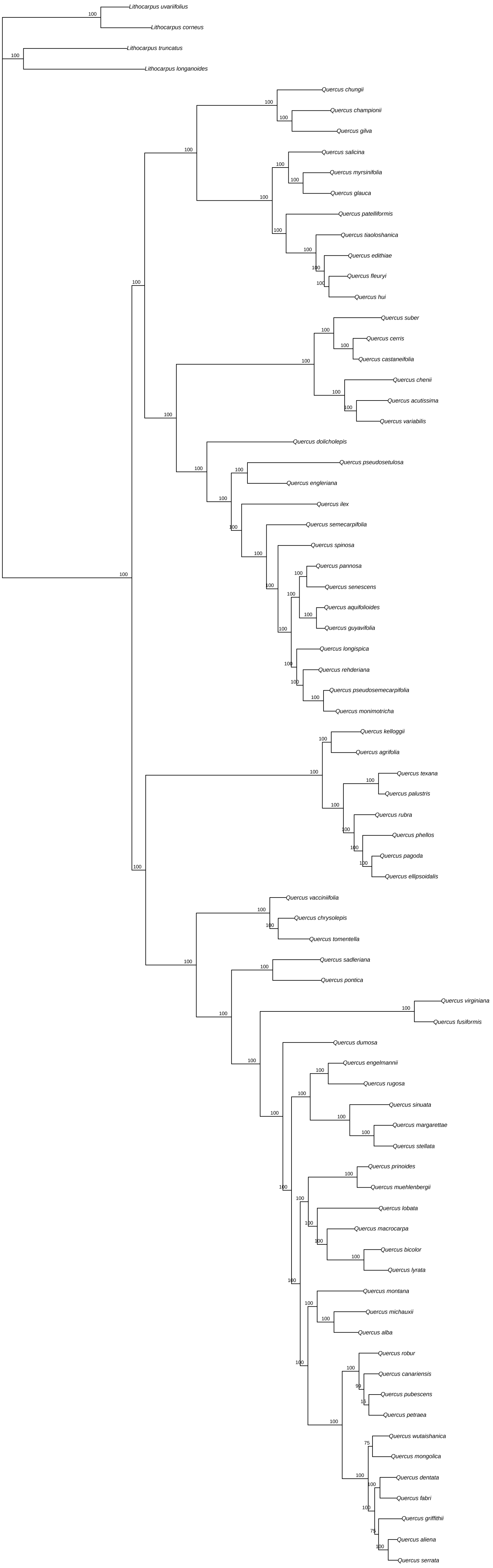

### Fig. S2b

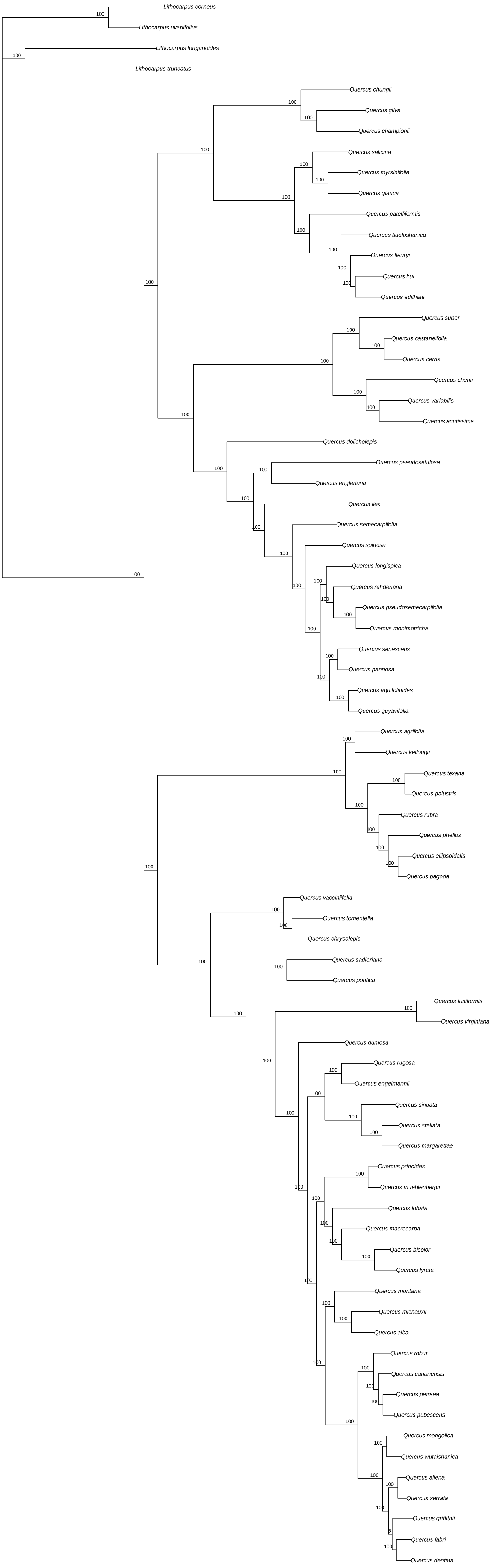

### Fig. S3a

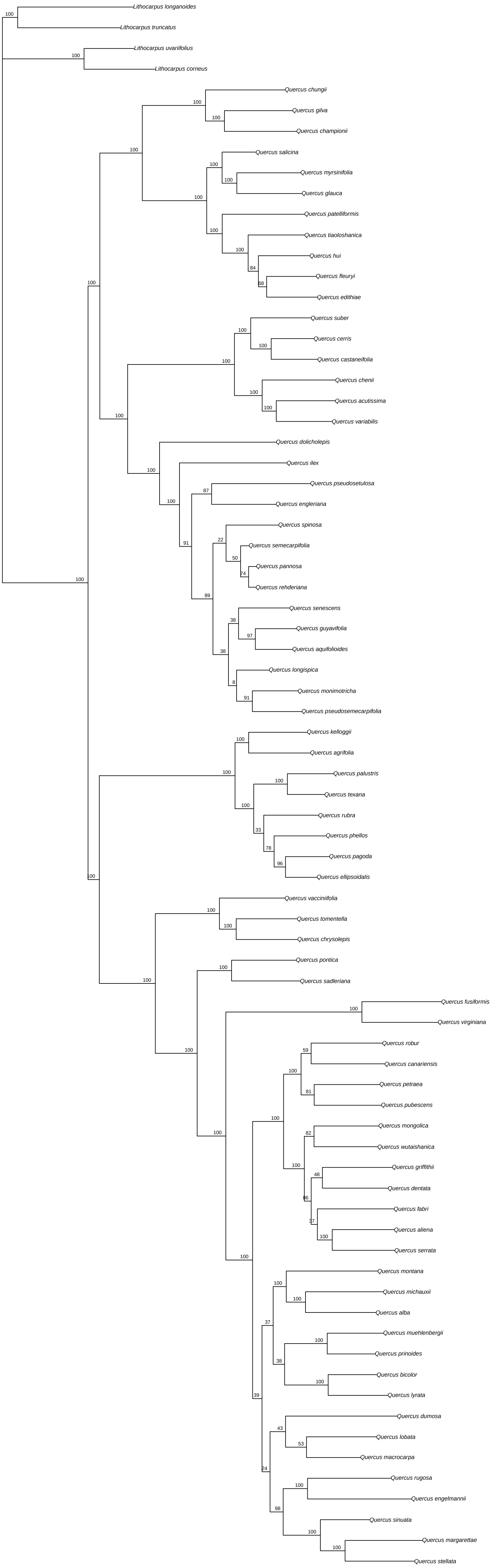

### Fig. S3b

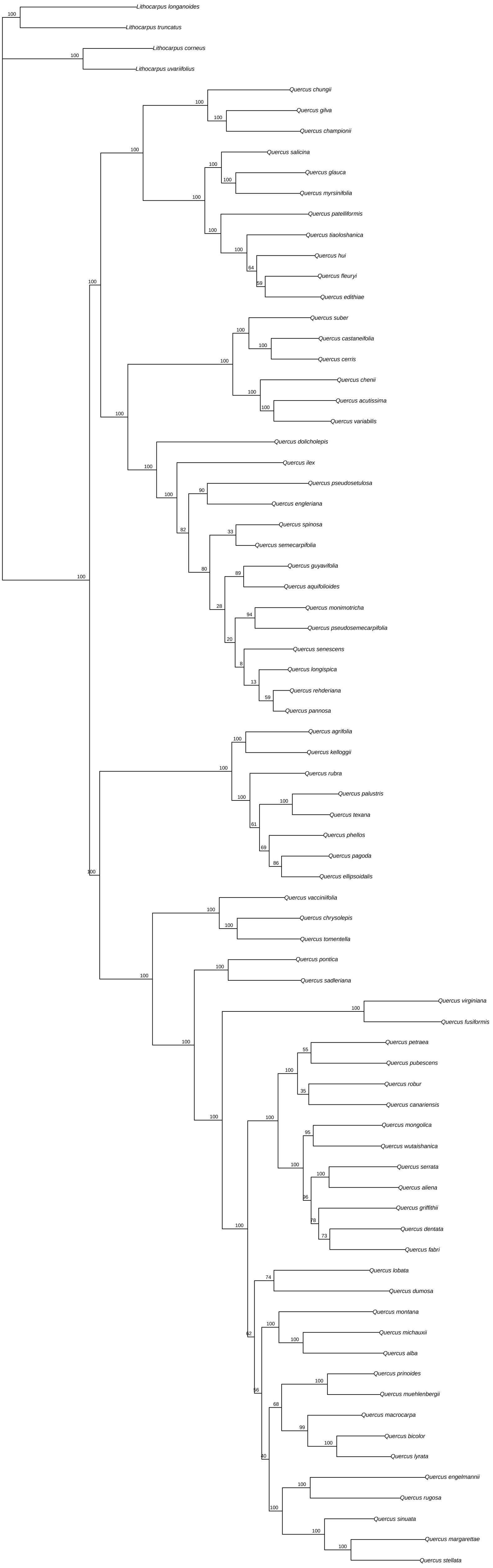

### Fig. S3c

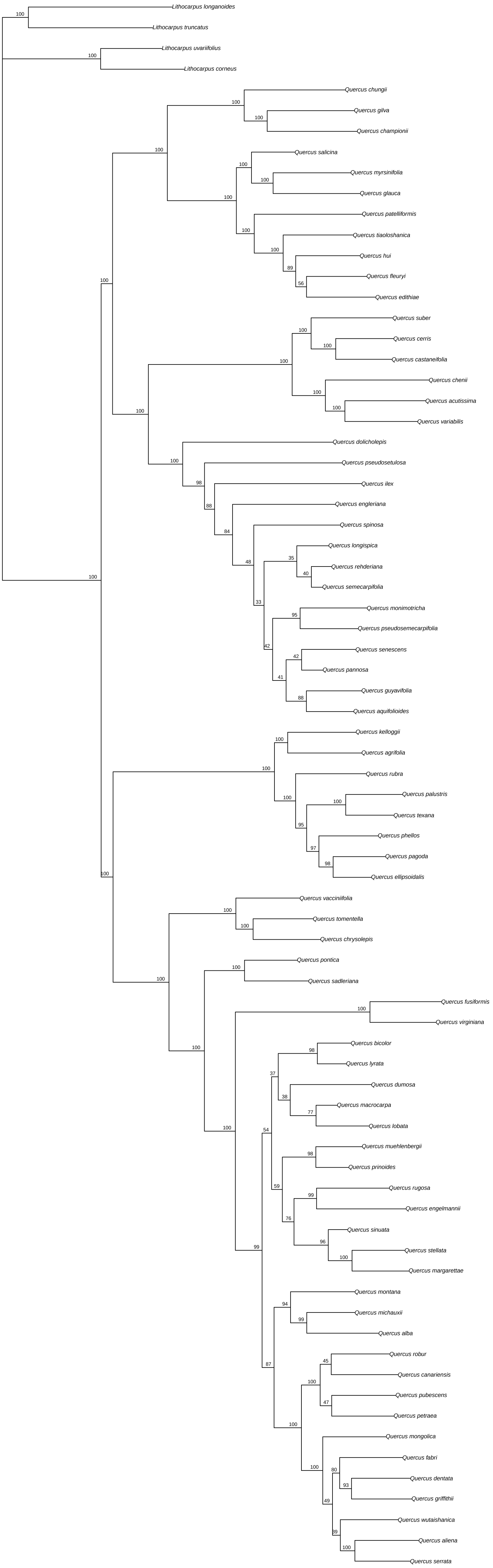

### Fig. S3d

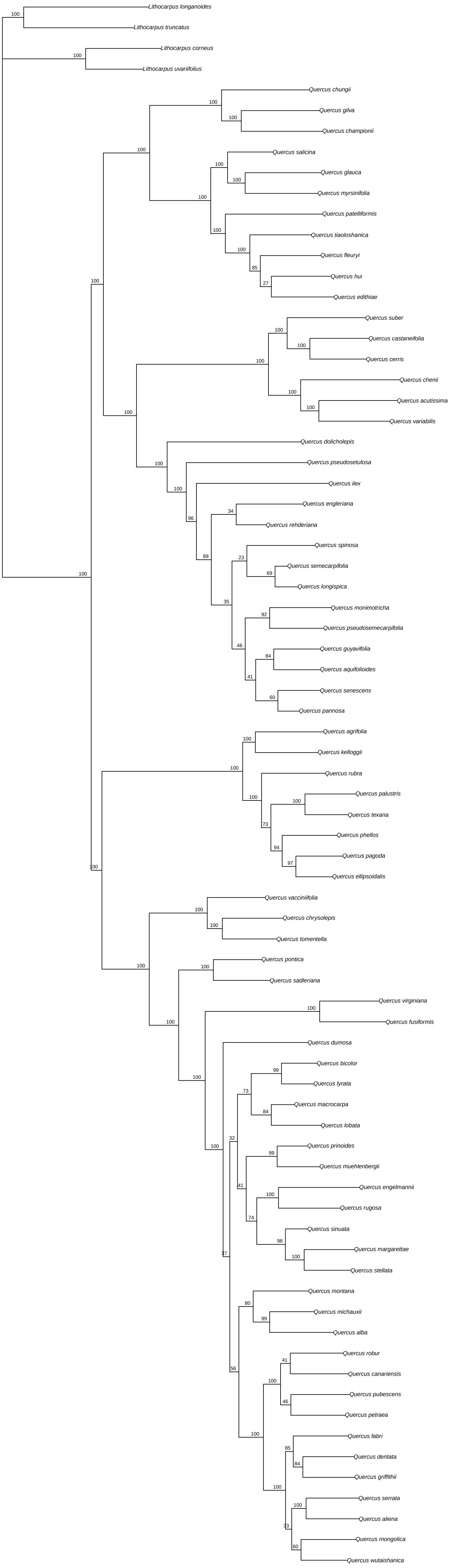

### Fig. S4

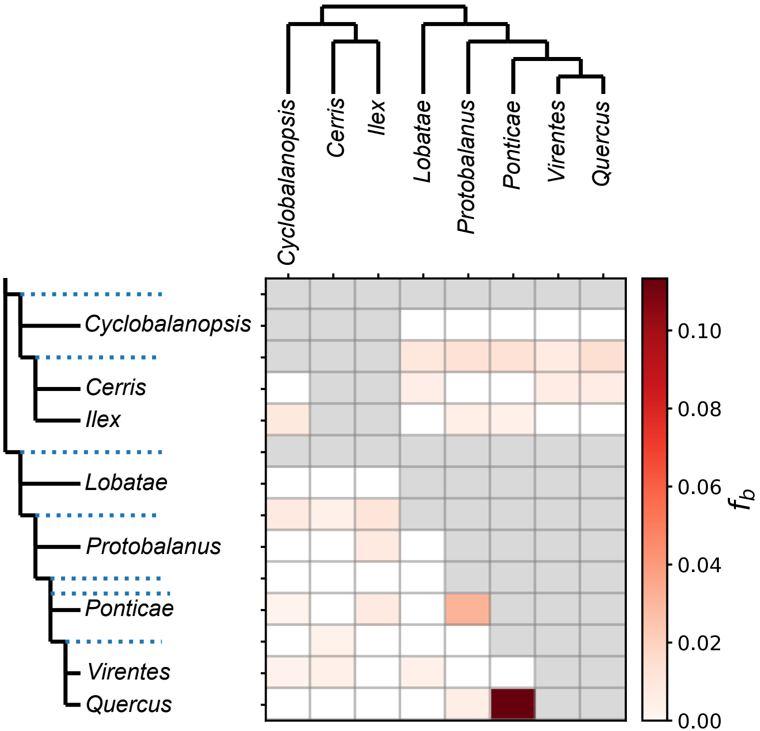

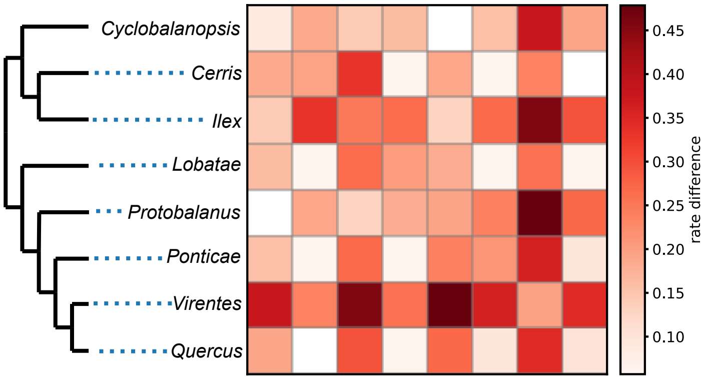

### Fig. S5

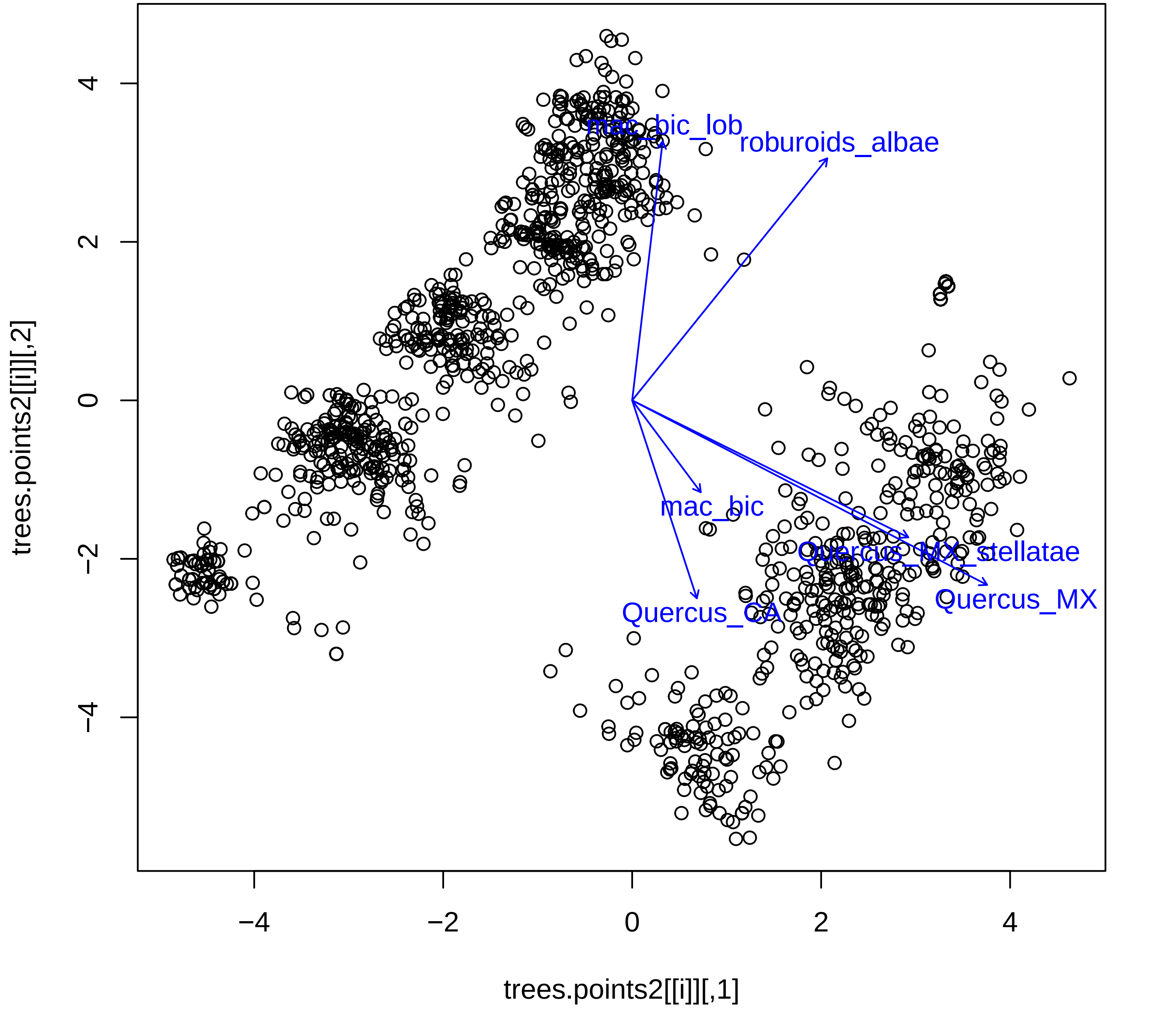
